# Structure of the E3 ligase CRL2-ZYG11B with substrates reveals the molecular basis for N-degron recognition and ubiquitination

**DOI:** 10.1101/2024.06.24.600508

**Authors:** Xi Liu, Yang Li, Lennice Castro, Zanlin Yu, Yifan Cheng, Matthew D. Daugherty, John D. Gross

## Abstract

ZYG11B is a substrate specificity factor for Cullin-RING ubiquitin ligase (CRL2) involved in many biological processes, including Gly/N-degron pathways. Yet how the binding of ZYG11B with CRL2 is coupled to substrate recognition and ubiquitination is unknown. We present the Cryo-EM structures of the CRL2-ZYG11B holoenzyme alone and in complex with a Gly/N-peptide from the inflammasome-forming pathogen sensor NLRP1. The structures indicate ZYG11B folds into a Leucine-Rich Repeat followed by two armadillo repeat domains that promote assembly with CRL2 and recognition of NLRP1 Gly/N-degron. ZYG11B promotes activation of the NLRP1 inflammasome through recognition and subsequent ubiquitination of the NLRP1 Gly/N-degron revealed by viral protease cleavage. Our structural and functional data indicate that blocking ZYG11B recognition of the NLRP1 Gly/N-degron inhibits NLRP1 inflammasome activation by a viral protease. Overall, we show how the CRL2-ZYG11B E3 ligase complex recognizes Gly/N-degron substrates, including those that are involved in viral protease-mediated activation of the NLRP1 inflammasome.

## INTRODUCTION

Members of the cullin-RING superfamily (CRLs) are modular protein complexes consisting of a substrate receptor, a cullin backbone and RING-box (Rbx) protein that binds coenzymes for ubiquitin transfer^1^. CRL2 is comprised of cullin-2(CUL2), Rbx1, and the adaptor heterodimer Elongin-B and Elongin-C (EloBC), which binds one of several substrate receptors^1^. ZYG11B and ZER1 are substrate receptors that directly bind proteins containing Gly/N degrons to recruit them to CRL2, forming the ubiquitin ligases complex CRL2-ZYG11B and CRL2-ZER1, respectively ^2^. As such, ZYG11B and ZER1 play a key role in protein quality control by promoting the degradation of Gly/N fragments formed during apoptosis and membrane proteins that fail to undergo myristoylation of N-terminal glycine^2^. However, how ZYG11B and ZER1 mediate substrate ubiquitination is unclear.

ZYG11B and ZER1 contain multiple domains of unknown function. Crystal structures of C-terminal ARM repeats of ZYG11B and ZER1 covalently fused to different Gly/N degrons were reported^3,4^. These studies revealed that a four-residue Gly/N degron binds to an N-terminal Glycine pocket (Gly-pocket). However, biochemical studies indicated that ZYG11B or ZER1 interact with their substrates using several different regions, which are not limited to the C-terminal ARM^3,4,5,6^. In addition to the Gly/N degradation pathway, ZYG11B has been implicated in the ubiquitination of substrates such as Cyclin B1, which do not contain Gly/N residues^5^. Structures of full-length ZYG11B and ZER1 are necessary to provide insight into the recognition of diverse substrates.

Human NLRP1 (NACHT, LRR, and PYD domains containing protein 1) has recently been characterized as a substrate of the ubiquitin CRL2 E3 ligase containing a Gly/N degron substrate receptor^7^ . NLRP1 is an effector-triggered immune (ETI) sensor of pathogen-encoded proteases, wherein cleavage of NLRP1 in mice and humans by specific bacterial or viral proteases results in inflammasome formation, leading to pyroptotic cell death and release of pro-inflammatory cytokines such as IL-1b and IL-18^7–10^. The mechanism by which this occurs, termed functional degradation, depends on the ’Function-to-find’ (FIIND) domain in NLRP1, which undergoes constitutive self-cleavage such that the N-terminal domains and the C-terminal caspase recruitment domain (CARD) remain noncovalently associated in an autoinhibited complex^11–16^. Following cleavage of a large ’tripwire loop’ in the N-terminus of NLRP1 by a pathogen-encoded protease, CRL2 recognizes and ubiquitinates the Gly/N degron at the neo-N-terminus of NLRP1. Ubiquitination of the neo-N-terminus of NLRP1 triggers proteasomal degradation and liberation of the CARD-containing C-terminus domain, which self-associates into a filamentous structure capable of activating caspase-1, leading to the release of inflammatory cytokines (e.g., IL-1b and IL-18) and pyroptotic cell death^11,12,17^. A similar activation mechanism of the related CARD8 inflammasome by virus cleavage has also been described^16,18–22^.

Evidence that NLRP1 is a substrate of CRL2 is provided by CRISPR-KO experiments: a double knockout of both ZYG11B and ZER1 reduces NLRP1 activation by the 3C protease from human rhinovirus (HRV)^7^. In support, treatment of cells with the pan-CRL inhibitor MLN4924 impairs HRV-mediated activation of NLRP1^7,23^. ZYG11B has also been implicated in other innate immune responses, such as potentiating the activation of cGAS during infection^24^. Consistent with its role at a host-pathogen interface, it has been proposed that ZYG11B is antagonized by viruses. For instance, degradation of ZYG11B is induced by HSV-1(herpes simplex virus type 1) infection^24^. In addition, the SARS-CoV-2 protein ORF10 interacts with ZYG11B and can suppress the innate immune response when overexpressed^25,26^. These data suggest that downregulation of the Gly/N degradation pathway may be a common viral strategy to evade immune surveillance. However, the way viruses specifically disrupt ZYG11B function remains to be determined. The above results suggest that CRL2-ZYG11B or CRL2-ZER1 are essential for NLRP1 activation after protease cleavage and that ORF10 may antagonize inflammasome activity.

Here we present the first Cryo-EM structure of CRL2-ZYG11B E3 ligase in its apo form as well as complexed with the Gly/N-degron of NLRP1 as a substrate. In the absence of substrate, ZYG11B forms a large protein interaction platform, making non-canonical interactions with the Cullin-backbone and EloBC. Substrate-bound complex structures reveal an extended surface for protein-protein interactions between ZYG11B and the Gly/N-degron of NLRP1 that goes far beyond the four-amino acid degron previously reported and defines the structural basis for substrate recognition by CRL2-ZYG11B. Consistent with these structural data, we show that ZYG11B knockout reduces NLRP1 inflammasome activation following cleavage by HRV 3C protease. Using functional studies and Cryo-EM, we also show that SARS-CoV2 ORF10 binds in a way that is mutually exclusive with the Gly/N degron of NLRP1 and prevents HRV 3C protease-mediated NLRP1 activation, suggesting a mechanism by which viruses could evade NLRP1-mediated sensing. Finally, our biochemical studies suggest a separate surface of ZYG11B is employed for binding substrates such as Cyclin B1, which does not have a Gly-N degron. Together, these observations suggest ZYG11B forms a large protein interaction platform to promote Gly/N-dependent and Gly/N-independent ubiquitination of substrates.

## RESULTS

### Structure of the ZYG11B-EloBC complex

To understand ZYG11B substrate recognition, we first determined the structure of the substrate receptor module ZYG11B in complex with EloBC (ZYG11B-EloBC) by single-particle Cryo-EM to a resolution of 4Å (**Extended Data** Fig. 1). ZYG11B binds to Elongin B and Elongin-C with 1:1:1 (single copy) stoichiometry. It contains a 3-helix motif bound in the prototypical CUL2 specificity factor von Hippel-Lindau protein (VHL) termed the VHL box, followed by Leucine Rich-Repeat (LRR) domain and an Armadillo Repeat (ARM) domain (**Fig. 1A**) ^27^. The full-length ZYG11B structure revealed that the VHL motif (or BC box) mediated the interaction between ZYG11B and Elongin-C. The LRR domain contains ten α/β horseshoe repeats forming a semicircular shape (**Fig. 1B**). The parallel β-strands form the concave face buried on the inside of the horseshoe, whereas the α-helical repeats form the convex face exposed to the solvent. Contacts between the LRR domain and ARM domain are mediated by intramolecular interactions (**Fig. 1C, Extended Data** Fig. 2A). The ARM domain harbors nine ARM repeats, but unlike most proteins with ARM domains, there is a long linker between ARM7 and ARM8 (635-652 a.a.). This linker insertion disrupts the tendency of the ARM repeat unit to fold together into a superhelix^28,29^. Instead, it breaks the ARM domain into two subdomains, which we term AD1 and AD2, respectively. The linker insertion enables the inversion of the AD2 domain so that it can fold back onto Elongin-B to stabilize the substrate receptor complex (**Fig. 1B**). Structural alignment of our full-length ZYG11B with previous crystal structures of AD2 of ZYG11B showed that only a very small portion of the protein interacts with Gly/N-degron, suggesting there could be additional protein-protein interactions between ZYG11B with substrate or the cullin backbone (**Fig. 1C**). In support, there is a large surface of conserved residues in ZYG11B from AD1, AD2 and the LRR, suggesting multiple protein interaction surfaces (Extended Data Figure 3).

**Fig. 1.**
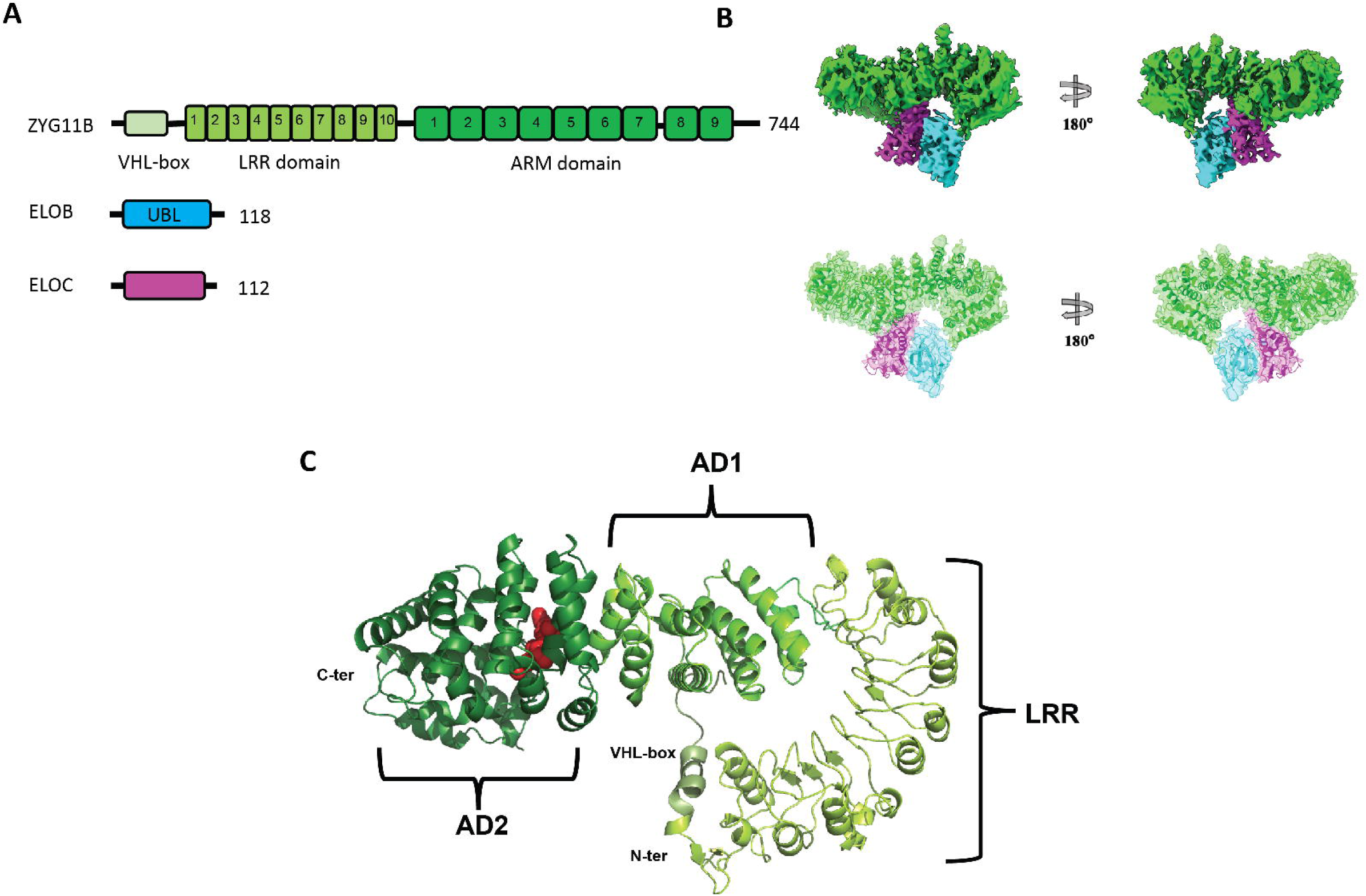
Structure of full-length ZYG11B in complex with Elongin-B and Elongin-C. **A** Schematic depicting the domain organization of ZYG11B, Elongin-B, and Elongin-C. **B**, Cryo-EM map (top) and model fitting into the maps of full-length ZYG11B in complex with Elongin-B/C complex (bottom). **C**, Cartoon model of ZYG11B displaying the VHL(BC-box), LRR, and ARM domains. The ARM domain is separated by the long linker between ARM7 and ARM8 and can be divided into two subdomains, AD1 and AD2. A small four-residue peptide was shown in the red sphere taken from the crystal structure of AD2 fused to the GFLH peptide (PDB: 7EP1).

### Structure of the CRL2-ZYG11B ubiquitin ligase

To address how the substrate receptor engages the cullin backbone, we determined the structure of the CRL2-ZYG11B complex. The structure of the CRL2-ZYG11B ubiquitin ligase was determined by single-particle Cryo-EM to a resolution of 3.8 Å (**Fig. 2A, Extended Data** Fig. 4 **and Extended Data** Fig. 5). This CRL2-ZYG11B E3 ligase forms a pentamer with single-copy stoichiometry (**Fig. 2B**). There are no significant structural rearrangements in ZYG11B-EloBC upon binding CUL2. We observe interactions between ZYG11B, EloBC, and CUL2 that have been reported in other substrate receptors for CRL2. For example, interfaces that mediate interactions between ZYG11B and CRL2 are present in the von Hippel-Lindau (VHL) factor, the prototypic substrate receptor for CRL2 (**Fig. 2C**) ^27,30^. The VHL box contains two alpha helices that are required for interaction with the EloBC heterodimer and CUL2, termed the BC box and the Cul2-box, respectively ^27,30,31^. In ZYG11B, the VHL box (residues L18 and C22) forms a hydrophobic interaction with residues Y76, L103, and L110 of Elongin-C, whereas residues in the CUL2 box of ZYG11B (P51, V54, F50) interact with CUL2 (residues Y107, T110) (**Fig. 2C**).

**Fig. 2.**
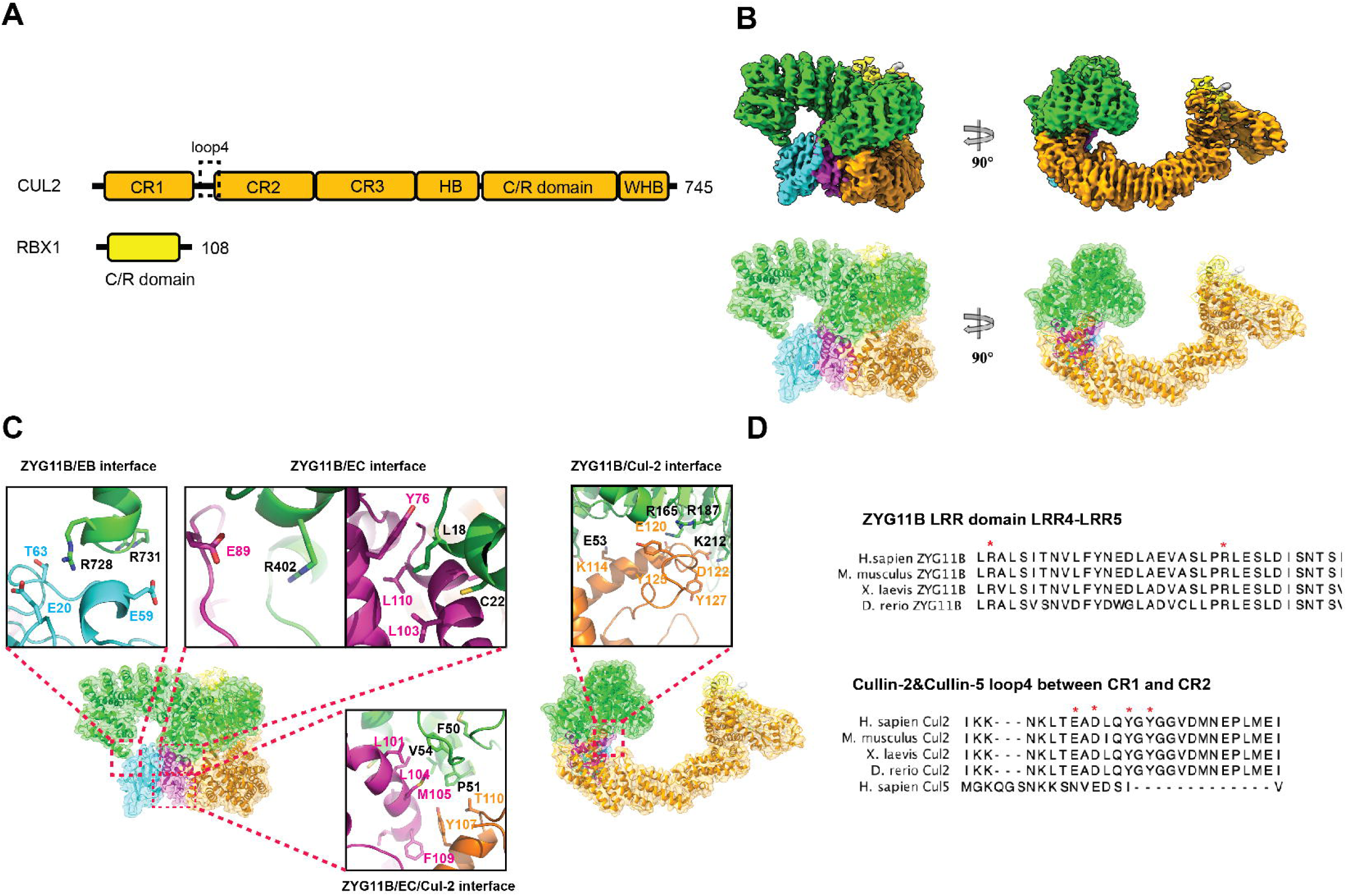
Structure of CRL-2 ZYG11B E3 ligase complex. **A** Schematic depicting the domain organization of CUL2 and Rbx1. **B**, Cryo-EM map (top) and model fitting into the map of the CRL2 ZYG11B E3 complex (bottom). **C**, Detailed interaction within the CRL2 ZYG11B E3 ligase complex assembly: the interface between ZYG11B CTD to Elongin-B (left); the interface between ZYG11B and Elongin-C; and three-way interface between ZYG11B, Elongin-C and CUL2 NTD (middle), a atypical interface between ZYG11B LRR with CUL2 loop4 (right). **D**, Sequence alignment of LRR4-LRR5 of ZYG11B among species (top), sequence alignment of CUL2 loop4 region between cullin repeat 1 (CR1) and CR2 among different species in comparison with the same region of CUL5 (bottom).

In contrast to the prototypical CRL2-VHL E3 ligase, we observe several unique interactions within the substrate receptor module and its complex with CUL2. For example, Elongin-B (E59) contacts ZYG11B (R728 and R731 of AD2), which may help stabilize the interaction between ZYG11B and the EloBC heterodimer (**Fig. 2C**). Additionally, loop 4 between cullin-Repeat 1 (CR1) and CR2 of CUL2 interacts with the LRR domain of ZYG11B (**Figs. 2A and 2C**, right). A similar binding mode was reported for *X. laevis* LRR1, a substrate receptor for CRL2 that promotes the termination of DNA replication^32^. Sequence conservation of loop 4, the observation that it interacts with CUL2 in our CRL2-ZYG11B structure, and its role in the previously reported structure of CRL2-LRR1 all suggest it functions in cullin selection (**Fig. 2D**).

### ZYG11B is required for activation of NLRP1

Prior studies indicate that ZYG11B and ZER1 are required for Gly/N degradation^2^. Double knockout of ZYG11B and ZER1 impaired the ability of HRV 3C protease to activate NLRP1 through the Gly/N degradation pathway^7^. To determine if ZYG11B is required for activation of NLRP1, we performed inflammasome reconstitution assays in cells where ZYG11B was knocked out using CRISPR/Cas9 (**Fig. 3**). HEK293 cells or those containing knockout of ZYG11B or a non-targeting control, were transfected with pro-IL-1b, NLRP1-TEV and TEV or human rhinovirus (HRV) 3C protease and the formation of mature IL-1b (p17) was monitored by western-blot (**Fig. 3A**). Cleavage of NLRP1-TEV by both TEV or HRV 3C protease led to inflammasome activation, as evidenced by maturation of IL-1b, consistent with prior studies^7,12,33^. TEV protease was able to activate NLRP1 in non-silencing control and in cells where ZYG11B was knocked out. In contrast, knockout of ZYG11B reduced activation of NLRP1 by HRV 3C protease compared to non-targeting control (**Fig. 3B, C**). We conclude that ZYG11B is required for activation of the NLRP1 inflammasome, but an additional E3 ligase may be required to promote inflammasome activation following TEV protease cleavage of NLRP1.

**Fig 3.**
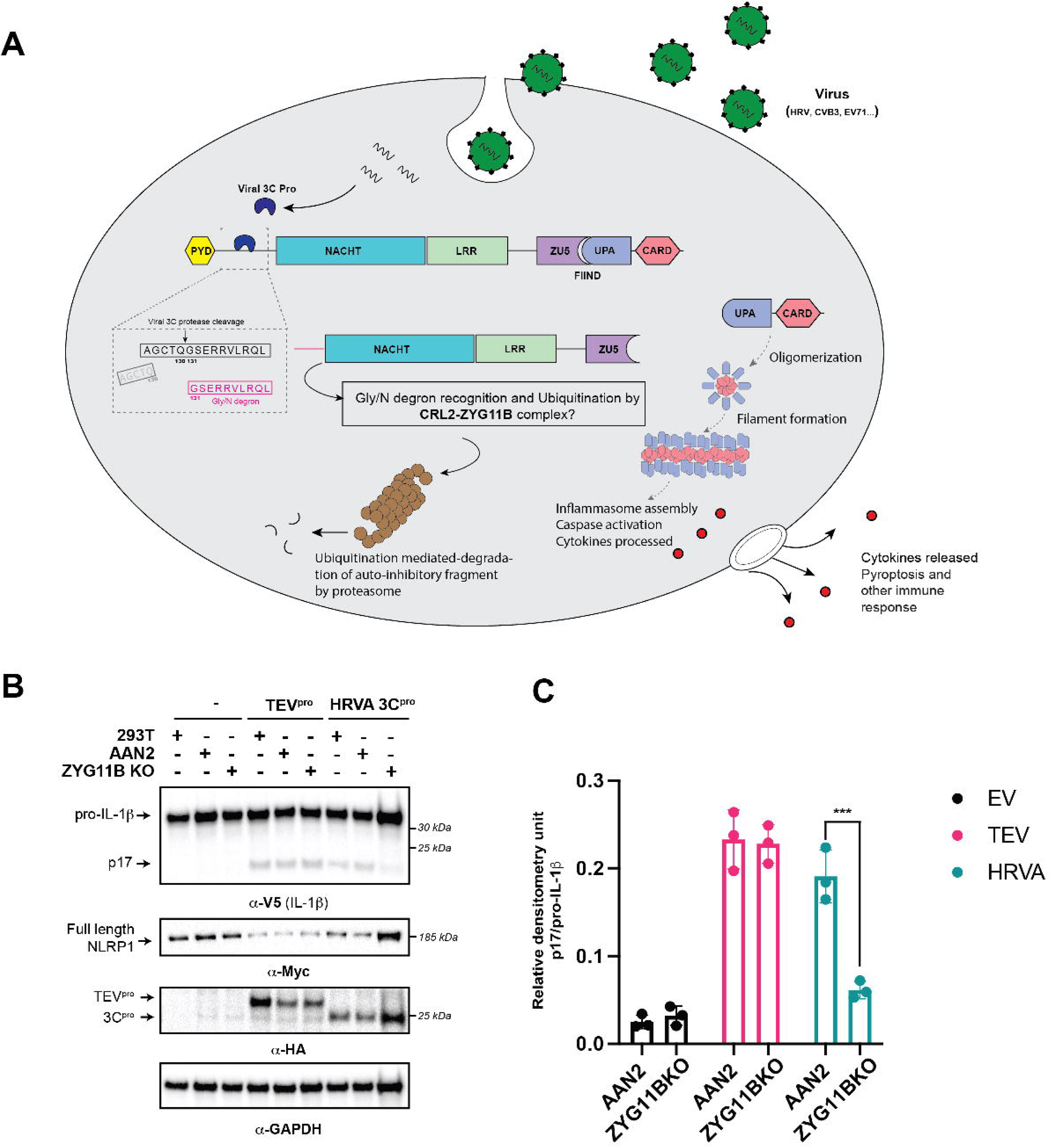
ZYG11B is required for activation of NLRP1 by human rhinovirus A (HRVA) 3C protease. **A** schematic showing the mechanism of activation of NLRP1. **B**, Western blot to assay inflammasome activation by formation of IL-1b (p17) in HEK293T cells transfected with pro-IL1b, NLRP1, HRV 3C or TEV protease where the ZYG11B (ZYG11B KO) or a safe harbor locus (AAN2) where targeted with CRISPR-CAS9. C, quantification of relative densitometry unit of p17/pro-IL-1b in panel B.

### Biochemical mapping of interactions between ZYG11B, NLRP1 and CRL2

We next asked if ZYG11B directly bound the NLRP1 Gly/N-degron produced by HRV 3C protease using MBP pulldown assays with recombinant purified components. Peptides containing 10 (Degron-10) and 20 (Degron-20) residues derived from 3C protease-cleaved NLRP1 were fused to the N terminus of MBP protein. Both the E3 holoenzyme (CRL2-ZYG11B) and its substrate receptor module ZYG11B-EloBC interacted with Degron-10 and Degron-20 MBP fusion protein (**Extended Data** Fig. 6A). In contrast, the MBP protein alone in the absence of the Gly/N-degron is not able to interact with the ZYG11B-EloBC complex. We conclude that the 10 and 20 amino acid peptides containing the Gly/N degron of NLRP1 are capable of directly binding ZYG11B.

Though EloBC can also bind Cullin-5(CUL-5), only CUL2 was detected in a genetic screen, identifying ZYG11B as an essential factor for Gly/N degradation^2^. We hypothesized that ZYG11B specifically binds CUL2. To test this possibility, we performed MBP pull-down assays using the Degron-10 and Degron 20 MBP fusion proteins to monitor the assembly of CRL2 or CRL5 with the substrate receptor complex ZYG11B-EloBC. Compared to the CUL2/Rbx1 complex, ZYG11B-EloBC could not form a stable complex with CUL5/Rbx2 (**Extended Data** Fig. 6A). These results suggest ZYG11B specifically binds CUL2 over CUL5, with loop 4 acting as a specificity determinant as suggested by structural studies (**Fig. 2C, D**).

Prior studies reported Cyclin B1 is a substrate of ZYG11B in vivo, yet Cyclin B1 was not shown to have a Gly/N degron^5^. We performed pull-down assays with recombinant purified Cyclin B1 and found it could interact with the ZYG11B-EloBC substrate receptor simultaneously with the Gly/N degron peptides from NLRP1 (**Extended Data** Fig. 6A). Accordingly, we hypothesized ZYG11B uses separate surfaces for interaction with Cyclin B1 and Gly/N degrons. To test this possibility, we generated a chimeric protein fusing the Gly/N degron of NLRP1 to the N-terminus of Cyclin B1. The purified chimeric protein copurified with ZYG11B-EloBC by size-exclusion chromatography, indicative of a stable complex and consistent with the notion that ZYG11B harbors separate binding sites for Cyclin B1 and Gly/N degrons (**Extended Data** Fig. 6B).

Next, we conducted Fluorescent Polarization assays to compare the binding of the known substrate and NLRP1 Gly/N degron to ZYG11B. Our findings reveal that the NLRP1 Gly/Degron peptide exhibits a binding affinity similar to that of peptides with high binding affinity in previous studies ^3^(**Extended data Fig. 6C**).

### ZYG11B recognizes the NLRP1 Gly/N-degron through AD1 and AD2 domains

We used information from the biochemical mapping experiments above to determine the structural basis of the NLRP1 Gly/N degron and Cyclin B1 binding to the CRL2-ZYG11B by determining the Cryo-EM structure of the E3 holoenzyme bound to an NLRP1 Gly/N degron covalently linked to Cyclin B1 (**Fig. 4**). Due to flexibility of Cyclin B1, we did not obtain high resolution of Cyclin B1 interface (**Extended Data** Fig. 4); however, the density of the Gly/N degron of human NLRP1 was resolved to 4.17 Å resolution (**Extended Data** Fig. 4 **and Extended Data** Fig. 5). Compared to CRL2-ZYG11B in the absence of substrate, the density for NRLP1 bound to the E3 ligase is clear (**Fig. 4B**). The Gly/N degron peptide of NLRP1 extends from the ARM domain (AD2) to the LRR domain. ZYG11B mainly uses three interfaces recognizing the Gly/N-degron of NLRP1, which we refer to as the Glycine pocket (Gly-pocket) that is buried in AD2, Extended Surface I, and Extended Surface II (**Fig. 4B**). The Gly-pocket forms interaction with the Gly/N degron NLRP1 in a manner consistent with prior structural studies^3^. W522 of ZYG11B forms a hydrophobic interaction with the main chain of the first residue of the Gly/N degron (NLRP1 G131, numbered as G1); D526 and N567 of ZYG11B form a hydrogen bond with the exposed protonated amino group of G1; and E570 of ZYG11B uses water-mediated hydrogen bond to interact with G1 of the Gly/N-degron. Mutation of residues in the Gly-pocket abolishes the binding between ZYG11B and NLRP1 degron (**Fig. 4C**). In contrast, the Extended Surface I and II have not been reported in prior studies due to lack of structural information on ZYG11B extending beyond AD2. At Extended Surface I, K515 E644 and K560 form a hydrogen bond and a salt bridge with NLRP1 Gly/N-degron residues R5, R4, and E3 respectively whereas Y685 interacts with R4 through a hydrogen bond (**Fig. 4B**). Compared to the Gly-pocket, mutation of Extended Surface I residues results in a moderate decrease in binding between ZYG11B and NLRP1(**Figs 4C and 4D**). The Gly/N-degron peptide extends further into the cavity of Extended Surface II, where F516 forms a cation-π interaction with R8 of the Gly/N-degron. Notably, R466 of ZYG11B uses its side guanidinium group to capture two individual carboxyl groups of the main chain of the NLRP Gly/N-degron through electrostatic interactions (**Fig. 4B**). The R466A mutation causes a relatively strong reduction in ZYG11B binding to the NLRP1 N degron. Mutation of residues in NLRP1 that contact ZYG11B in our structure also reduces the interaction (**Extended Data** Fig. 7). For example, the E133K mutation of NLRP1 (residue E3 of the Gly/N degron) decreases the interaction between ZYG11B, indicating that the P3’ position of the rhinovirus 3C protease site of NLRP1 determines the specificity of substrate of N glycine degron (**Extended Data** Fig. 7). This result agrees with previous direct binding data showing that residues at P3’ position could change binding affinity of ZYG11B CTD to Gly/N-degron ^3^. Our results suggest that the degron of NLRP1 recognized by ZYG11B extends well beyond the two amino acids contacted by the Gly-pocket ^3^. We conclude that ZYG11B binding to the NLRP1 Gly/N degron not only requires the N-terminal Glycine residue but also a longer peptide going through the channel formed by Extended Surface I and II.

**Fig. 4.**
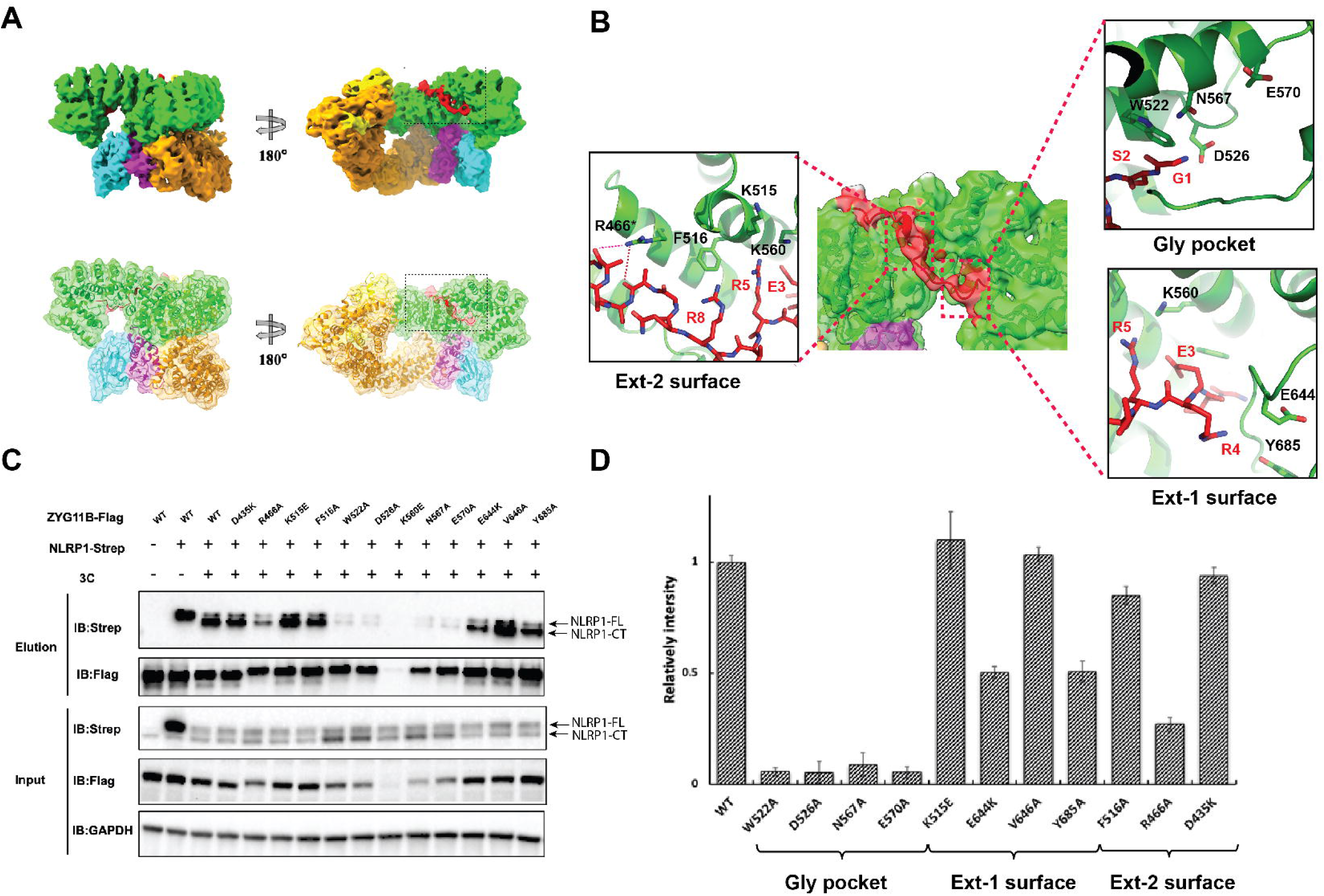
ZYG11B recognition of the NLRP1 Gly/N degron. **A**, Cryo-EM map (top) and model fitting into the maps (bottom) of CRL-2 ZYG11B E3 complex binding to NLRP1 Gly/N degron. **B**, Detailed interactions between ZYG11B and Gly/N degron of NLRP1. Three interfaces mediate the interaction: the previously observed glycine recognition pocket (gly pocket), the extended surface I (Ext-1), and the extended surface II (Ext-2) spanned by AD1 and AD2 of ZYG11B. The Gly/N degron of NLRP1 generated by HRV protease cleavage begins at Glycine-131, labeled as G1 in the figure. **C**, Co-IP analysis (with anti-Flag) and western blot analysis (with anti-Flag or anti-Strep) of HEK 293T cells transfected with plasmids encoding Strep-tagged NLRP1 and Flag-tagged WT ZYG11B and ZYG11B mutants (D435K, R466A, K515E, F516A, W522A, D526A, K560E, N567A, E570A, E644K, V646A and Y685A) for 48 hr. **D**, Quantification of intensities in panel C. The relative intensities were normalized to WT NLRP1 (digested NLRP1 by 3C protease in elution/digested NLRP1 by 3C protease in input) with residues categorized by surfaces depicted in panel B.

### ORF10 binds to ZYG11B mutually exclusive with NLRP1 and can inhibit inflammasome activation

We next asked if Gly/N degrons with alpha-helical secondary structures engage ZYG11B using similar binding modes as the linear motifs characterized to date^3,4,6^. Though the role of ORF10 for SARS-CoV-2 infection is still under debate, ORF10 is predicted to be alpha-helical, so it could be a valuable tool to investigate the specificity of ZYG11B and whether virally encoded peptides could inhibit inflammasome activation^25,26,34–41^. Accordingly, we determined the Cryo-EM structure of the substrate receptor module ZYG11B-EloBC with a Gly/N terminal peptide from ORF10 (**Fig.5A, Extended Data** Fig. 1 **and Extended Data** Fig. 5). The overall structure of the ZYG11B-EloBC complex is very similar to that we observe bound to CUL2/RBX1 in the CRL2-ZYG11B complex. The first 16 residues of ORF10 are located at the Gly-pocket and extended surfaces, similar to NLRP1 degron. The N-terminal Glycine of ORF10 was captured by W522, D526, N567 and E570 at Glycine pocket (**Fig. 5B**). Compared to NLRP1, there are fewer interactions observed at Extended Surface I: the N4 residue of ORF10 forms a hydrogen bond with E644. The phenolic hydroxyl group of Y2 of ORF10 forms a hydrogen bond with R649. In contrast to NLRP1, ORF10 uses a hydrophobic surface to interact with ZYG11B at extended surface II: residues from a short helix of ORF10 (F7, P10 and I12) interact with ZYG11B Extended Surface I (I474 and F516) by forming a hydrophobic interaction (**Fig. 5B**). Despite different binding modes at Extended Surface I and II, structural alignment of ORF10 and NLRP1 in complex with ZYG11B predicts mutually exclusive binding (**Fig. 5C**).

**Fig. 5.**
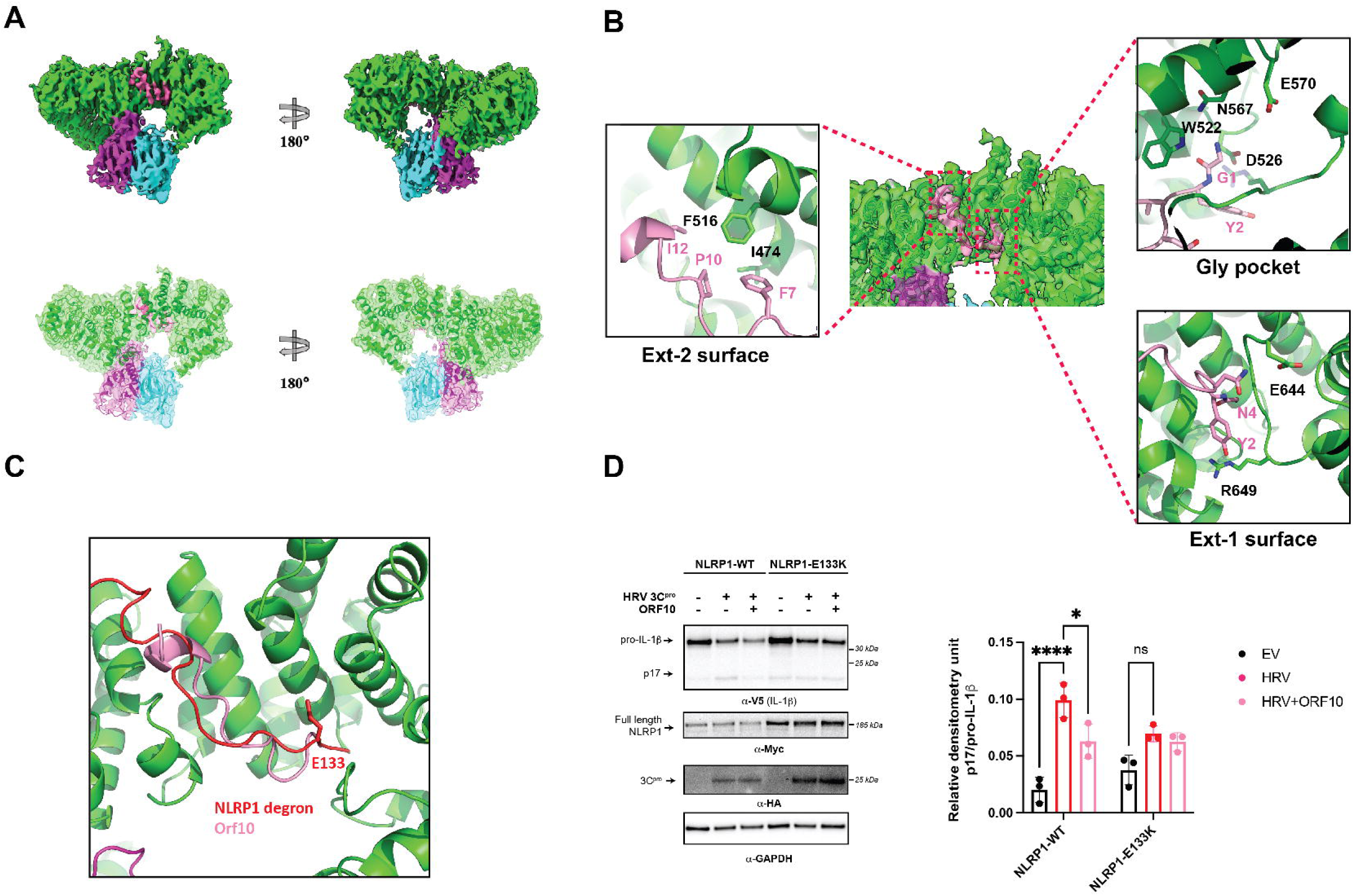
Structure of substrate receptor module ZYG11B-EloBC bound to SARS-CoV2 ORF10 **A**, Cryo-EM map (top) and model fitting into the maps (bottom) of ZYG11B EloBC complex binding to SARS-Cov2 ORF10(pink). **B**, Detailed interaction between ZYG11B and ORF10. The interfaces space three surfaces: the glycine recognition pocket (Gly pocket), extended surface I (Ext-1), and extended surface II (Ext-2). **C**, Structural alignment of NLRP1 degron (red) with ORF10(pink) on the binding surface of ZYG11B, predicting mutually exclusive interactions. **D**, Western blot of pro-IL1-β and p17 produced by HEK293T cells transfected with HRV3Cpro, ORF10, NLRP1, and its mutant as indicated (left)—quantification of intensities of p17 formation (right).

To determine whether ORF10 could functionally compete with NLRP1 degron binding to ZYG11B, we tested if overexpression of ORF10 would inhibit NLRP1 inflammasome activation following cleavage by HRV protease. As anticipated, the overexpression of ORF10 could inhibit the HRV 3C protease-mediated activation of NLRP1, evidenced by decreased maturation of IL-1b in the presence of ORF10 overexpression (**Fig. 5D**). In contrast to wild-type, the E133K mutation of NLRP1 decreased its binding to ZYG11B and reduced inflammasome activation by HRV protease (**Extended Data** Fig. 7, **Fig. 5D**). In this case, ORF10 expression did not have a measurable effect on NLRP1 inflammasome activation (**Fig. 5D)**. In conclusion, our structural and cell-based studies indicate SARS-CoV2 ORF10 could inhibit NLRP1 inflammasome activation through functional degradation by ZYG11B.

## DISCUSSION

In this study, we determined the structure of the E3 ligase CRL2-ZYG11B complex, as well as the complex between CRL2-ZYG11B and a substrate, the Gly/N degron produced when NLRP1 is cleaved by human rhinovirus (HRV) 3C protease. We further showed that the Gly/N degron receptor ZYG11B is required for activation of the NLRP1 inflammasome by HRV 3C protease. In addition, we determined the structure of SARS-CoV-2 ORF10 in complex with ZYG11B, which suggests effector-triggered inflammasome activation could be blocked by viral mimicry of Gly/N degron motifs **(Fig. 6)**. The implications of these findings for the mechanism of NLRP1 activation and inhibition will be discussed below.

**Fig. 6.**
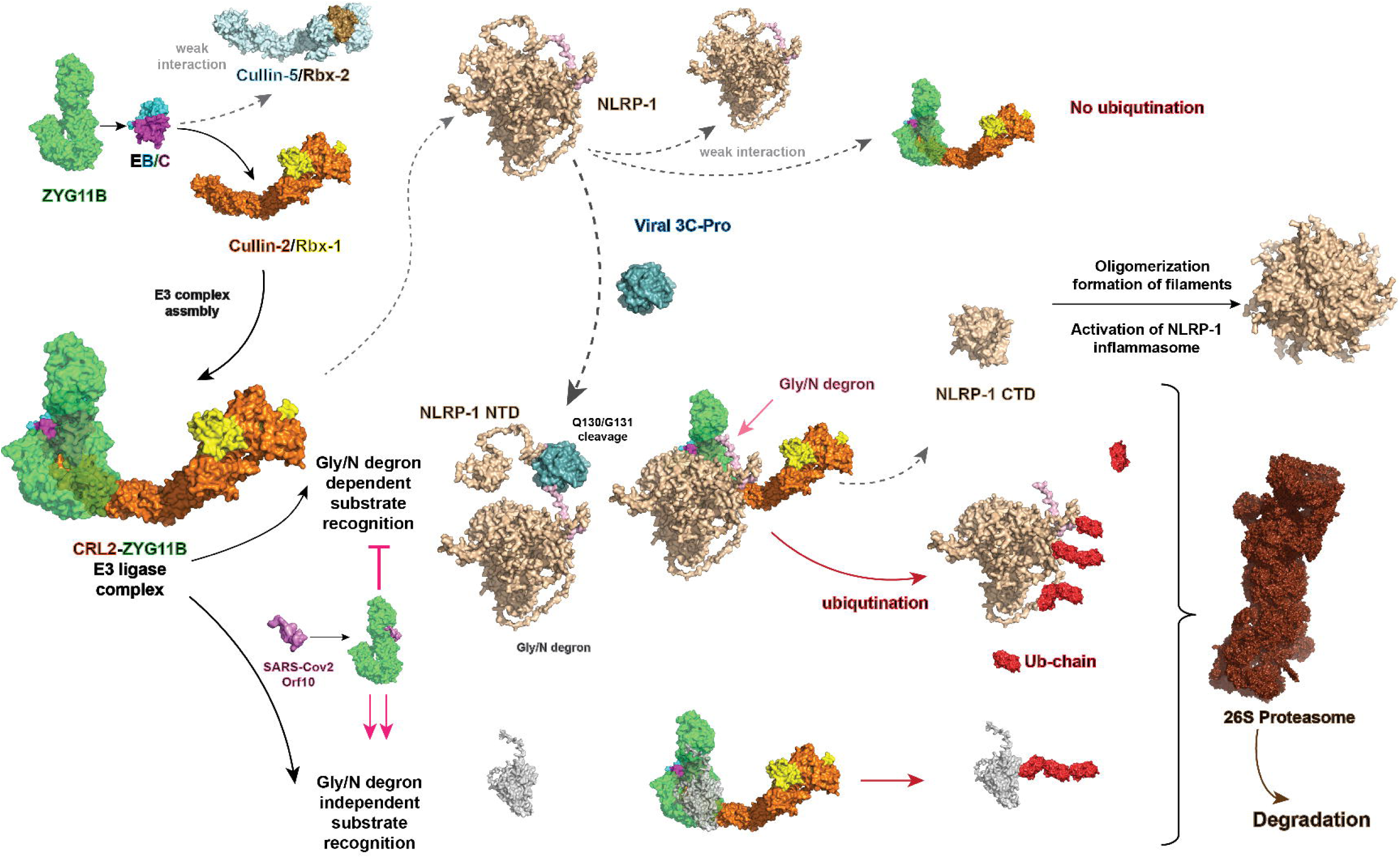
Proposed model of ZYG11B requirement for NLRP1 activation via sensing viral protease and the inhibition by SARS-CoV2 ORF10. ZYG11B specifically recruits Cullin-2/RBX1 to form the E3 ligase complex. In the absence of viral infection, ZYG11B exhibits minimal or no interaction with NLRP1. Upon viral infection, 3C or 3C-like proteases cleave NLRP1, exposing the Gly/N degron. This cleavage event facilitates the recognition of NLRP1 by ZYG11B, leading to its ubiquitination by the CRL-2 ZYG11B E3 ligase complex. The ubiquitination process results in the release of the auto-inhibitory N-terminal fragment from the C-terminal domain (CTD) of NLRP1. Consequently, the CTD of NLRP1 oligomerizes and forms filaments within cells, triggering the activation of the NLRP1 inflammasome and subsequent immune responses. SARS-CoV-2 ORF10 and other viral proteins can mimic the Gly/N degron of NLRP1, thereby inhibiting Gly/N degron-dependent substrate recognition and suppressing the immune response through targeted degradation.

ZYG11B is a large protein that folds into multiple domains a LRR flanked by two armadillo repeat domains that we refer to as AD1 and AD2. Prior crystallographic studies where four amino acids Gly/N degron are covalently fused to AD2 reveal how glycine interacts with the Gly-pocket. Using full-length ZYG11B, we show that an additional binding site is formed at the junction of AD1 and AD2, consisting of Extended-surface 1 and 2, which together enable recognition of Gly-N degrons containing around a dozen amino acids. While recognition of the N-terminal glycine is highly specific, neighboring residues can provide additional affinity through interactions of the backbone with residues within Extended Surface 1 and 2, respectively.

Prior genetic and biochemical studies indicate Cyclin B1 is a substrate for the CRL2-ZYG11B E3 ligase, yet Cyclin B1 does not possess a Gly/N degron^5^. Our biochemical studies reveal that Cyclin B1 and Gly/N degrons can bind ZYG11B simultaneously, suggesting separate surfaces of ZYG11B mediate the binding of Cyclin B1 and Gly/N degrons (**Extended Data** Figure 6 **and Fig 4**). These observations suggest: 1-ZYG11B acts as a substrate receptor for folded proteins such as Cyclin B1 independent of Gly/N degradation and 2-that the Cyclin B1 binding surface on ZYG11B might be utilized for some Gly/N substrates to provide additional affinity or positioning of substrate receptor lysines for ubiquitination by CRL2. Testing these possibilities will require additional functional studies with full-length proteins containing Gly/N degrons.

The composite interaction surface formed by the junction of AD1 and AD2 we describe in our structure may enable recognition of an extensive array of primary sequences and secondary structures near the short four amino acid Gly/N degron previously described ^3,4,6^. Consistent with the latter possibility, we observed that SARS-Cov-2 ORF10 binds to ZYG11B using the classic four amino acid degron flanked by an alpha-helix that binds the junction between AD1 and AD2, suggesting ORF10 binds mutually exclusive with Gly/N degrons (**Fig 5**). While the first four amino acids of ORF10 and NLRP1 Gly/N degrons bind the Gly-pocket, they extend into the junction between AD1 and AD2 using alpha-helical or extended secondary structures, respectively. The common binding surface of Gly/N degrons of NLRP1 and ORF10 on ZYG11B predict mutually exclusive interactions (**Fig 5C**). These observations are consistent with the detection of CRL2-ZYG11B as an interaction partner of ORF10 in a high-throughput protein interaction screen^25^.

We further find that ORF10 inhibits the activation of NLRP1 by HRV 3C protease (**Fig 5D**). While these data indicate that overexpression of the ORF10 protein can inhibit NLRP1 inflammasome activation, the role of ORF10 in SARS-CoV2 infection remains unclear. Several studies suggest mRNA is not detected in various cell lines infected with SARS-CoV2^34–36,38^. However, more recent RNA-seq data from 2070 samples from diverse human cells and tissues reveals the expression of ORF10 RNA^41^. Moreover, ORF10 is not required for viral replication in HEK293T cells expressing the ACE2 receptor^21^, but has been shown to be required for replication of SARS-CoV-2 in a hamster model in vivo^40^, and an intact ORF10 gene of SARS-CoV-2 was shown to be correlated with more severe COVID-19 in the human host--in contrast to missense and non-sense mutation within ORF10^41^. While our results do not resolve whether ORF10 is expressed during SARS-CoV-2 infection in vivo, it is tempting to speculate that the expression of ORF10 could reduce inflammasome activation during SARS-CoV-2 infection. Indeed, NLRP1 was recently shown to be a sensor of SARS-CoV-2 infection in lung epithelia via the detection of SARS-CoV-2 3CL protease cleavage of NLRP1 at position Q333 within the NACHT domain, which exposes a Gly/N degron^10^. Moreover, CARD8 is an NLRP1-like inflammasome with a mechanism of activation that proceeds through functional degradation, and SARS-CoV-2 3CL protease activates CARD8 and reveals a Gly/N degron^19^. Testing the possibility that viral proteins such as ORF10 antagonize Gly/N degradation through molecular mimicry as a mechanism to reduce inflammasome activation is a challenge for the future.

In recent years, targeted protein degradation (TPD) has emerged as a novel therapeutic approach, particularly for cancer treatment. The innovative concept behind proteolysis-targeting chimeras (PROTACs) involves simultaneously recruiting a protein of interest and an E3 ubiquitin ligase. While notable successes have been in targeting the CRL2 E3 ligase, the focus has primarily centered on VHL^42^.

Our research could also provide valuable structural insights necessary for the development of PROTACs targeting the CRL-ZYG11B E3 ligase to expand the repertoire of E3 ligases available for PROTAC design.

In summary, we have uncovered an unexpected plasticity in the recognition of substrates by the E3 ligase CRL2-ZYG11B. This E3 ligase is required for the functional degradation of NLRP1 during innate immunity and can recognize Gly/N degrons flanked by diverse sequences and secondary structures. Our work further suggests that inhibition of Gly/N degradation could be a viral strategy for suppressing innate immunity through molecular mimicry.

## Acknowledgments

Thanks are accorded to Dr. Stephen Elledge of Harvard Medical School for the gift of the ZYG11B CRISPR KO cell line and members of the Gross and Daugherty labs, Drs David Gordon and Nevan Krogan for sharing data prior to publication. This work was supported by grants from the Sandler Family Program for Breakthrough Biomedical Research and the Big Blue Sky Foundation ( to JDG) and grants from the National Institutes of Health (NIH) ( U54AI170792)(to JDG). This work was supported by grants from the National Institutes of Health (NIH) (R35 GM133633) and Burroughs Welcome Investigators in the Pathogenesis of Infectious Disease Program to MDD. LKC was supported by a Ford Foundation Predoctoral Fellowship and an HHMI Gilliam Fellowship for Advanced Study.

## Author contributions

XL reconstituted the CRL2-ZYG11B E3 ligase and binding partners and performed Cryo-EM structure determination with ZY. YL performed co-IP experiments. LC performed inflammasome activation assays. XL and JDG conceived the project. YC, MD, and JDG supervised the research. All authors contributed to the interpretation of the data and writing of the manuscript.

## Declaration of interests

The authors declare no competing interests.

## Figure Legends

**Extended Data Fig 1.** Structure determination of the ZYG11B-EloBC complex and ZYG11-EloBC bound to SARS-CoV2 ORF10 **a**, Purified ZYG11B-EloBC complex verified on SDS-PAGE by Coomassie blue staining. **b**, Representative cryo-EM micrograph. **c**, Representative 2D class averages. **d**, workflow for the structure determination of the ZYG11B-EloBC complex. **e**, workflow for the determination of the structure of ZYG11B-EloBC complex with SARS-CoV2 ORF10 protein.

**Extended Data Fig 2.** Close-up of interactions predicted to stabilize intra and intermolecular interactions of ZYG11B. **A**, interactions between the LRR and ARM domains of ZYG11B. **B**, the overlay of CUL5 and CUL2 showing differences in the conformation of loop-4. CUL2 was taken from the CRL2-ZYG11B complex determined in this study, and CUL5 from the Alphafold2 protein structure database.

**Extended Data Fig 3.** Surface conservation analysis of ZYG11B across species. The 3D structure of the ZYG11B surface is color-coded based on Consurf conservation scores, transitioning from variable (blue) to conserved (red), derived from sequence alignment of available sequences depicted on the right. Secondary structures coordinating with the sequence are illustrated and labeled.

**Extended Data Fig 4.** Structure determination of the CRL2-ZYG11B complex (**A-D)** and CRL2-ZYG11B bound to the NLRP1 Gly/N degron fused to Cyclin B1 (**E-G**).

**a**, Purified CRL2-ZYG11B complex verified on SDS-PAGE by Coomassie blue staining. **b**, Representative cryo-EM micrographs. **c**, Representative 2D class averages. **d**, Workflow for structure determination. **e**, representative cryo-EM micrograph of CRL2-ZYG11B bound to Cyclin_B1-NLRP1 Gly/N degron fusion peptide. **f**, Representative 2D class averages. **g**, workflow for structure determination.

**Extended Data Fig 5.** Representative Cryo-EM density maps of the CRL2-ZYG11B complex and Gly/N degron fitting to model.

**Extended Data Fig 6.** Biochemical characterization of NLRP1 Gly/N degron and Cyclin B1 binding to ZYG11B. **A**, Pull down assays of MBP-fused NLRP1 Gly/N degron with ZYG11B/EloBC complex together with CUL2/Rbx1, CUL5/Rbx2, and Cyclin B1. The first 10 or 20 residues of the NLRP1 Gly/N degron produced by HRV A 3C protease cleavage (residues 131-141 or 131-151) were fused to MBP, labeled Degron_10 and Degron_20 respectively. **B**, Size exclusion chromatography showing binding of Cyclin B1 with ZYG11B-EloBC. A major peak shift was observed when Cyclin B1 and ZYG11B-EloBC complex were incubated**. C**, fluorescence anisotropy binding of ZYG11B-EloBC to indicated fluorescein-labeled peptides (left) and fitted equilibrium dissociation constants (right).

**Extended Data Fig 7.** Co-Immunoprecipitation of ZYG11B with wild-type or indicated mutants of NLRP1. **A**, zoom in of interactions between ZYG11B and Gly/N degron of NLRP1. b, Flag-tagged ZYG11B and indicated variants of strep-tagged NLRP1 were cotransfected into HEK293T cells, subjected to immunoprecipitation with anti-flag resin, and assayed by western blot.

## Methods Details

### Protein expression and purification

The gene encoding human ZYG11B was codon-optimized for *E. coli* expression and amplified by PCR. The PCR products were cloned into a modified pRSF-Duet-SUMO vector. The recombinant plasmid contains a 10x His-SUMO tag at the N-terminal and could be removed by Ulp1 protease. Elongin-B(1-118) and Elongin-C(17-112) were cloned to the pCDF-Duet vector. These two plasmids were co-transformed into *E. coli* BL21(DE3) cells with Kanamycin and Streptomycin selection. The cells were cultured in Luria-Bertani (LB) medium at 37 °C until the value of OD600 reached 0.8-1.0, then the temperature was reduced to 16 °C. The protein complex expression was induced by 0.5mM isopropyl-D-1-thiogalactopyranoside (IPTG) for 12-16hrs culture. The *E. coli* cells containing the target protein were harvested by centrifugation at 4000 rpm for 15 min. The cells were resuspended in lysis buffer A composed of 50mM Tris-HCl pH 8.0, 500mM NaCl, and 20mM imidazole. The cells were lysed by sonication on cell, and insoluble cellular debris was removed by centrifugation at 14,500 rpm for 1hr. The supernatant was then flowed through a prepacked Ni-NTA column at 4°C. After loading the supernatant, the column was washed by washing buffer containing 50mM Tris-HCl pH 8.0, 500mM NaCl, and 40mM imidazole. The complex of SUMO-ZYG11B-EloBC was eluted by elution buffer (25 mM Tris-HCl pH 8.0 150mM NaCl and 500mM imidazole pH 8.0). The SUMO tag was cleaved by homemade Ulp1 protease with a molar ratio of 1:100 at 4 °C overnight. The sample was exchanged to Q buffer A (25mM Tris-HCl pH8.0 80mM NaCl and 0.5mM TCEP). Further purification was performed by anion exchange chromatography (HiTrap Q HP column, Cytiva) with a gradient of NaCl from 80 to 800mM to remove the SUMO tag and Ulp1 protease. Purified ZYG11B-EloBC complex was concentrated and injected into size exclusion chromatography column (Superose 6 Increase 10/300 GL, GE Healthcare) pre-equilibrated with Sizing buffer (20mM Hepes pH7.5 300mM NaCl and 1mM TCEP). The purified homogenous ZYG11B-EloBC complex protein was verified by SDS-PAGE, concentrated, and stored at -80 °C for later usage.

The human CUL2 and Rbx-1 complex was expressed using the Bac-to-Bac baculovirus expression system (Invitrogen). The genes were amplified by PCR from cDNA, with a twin-StrepII tag added to the C terminus of CUL2. The PCR products were cloned into pFast-Bac-Dual vector. The construct was transformed into bacterial DH10Bac cells to generate bacmid. The extracted bacmid was then transfected into *Spodoptera frugiperda* Sf9 insect cells to produce recombinant baculovirus. Low-titer viruses were harvested and then amplified by two more rounds. The amplified virus was used to infect 4 liters of Sf9 cells at a density of 2x 106 cells per milliliter. The cells were harvested after 60hrs infection by centrifugation at 4000rpm for 15mins and then resuspended in lysis buffer B composed of 50mM Tris-HCl, 500mM NaCl, 0.5mM EDTA,0.5mM TCEP and 1mM PMSF. After adding protease inhibitor cocktail, the cells were lysed by sonication and ultracentrifuged at 40,000 rpm for 1hr to remove cell debris. The supernatant was incubated with Strep-Tactin Sepharose (IBA) resin for 2hrs. The resin was then extensively washed with lysis buffer B, and CUL2/Rbx-1 complex was eluted using Strep Elution buffer (20 mM Hepes pH7.5 300mM NaCl 0.5mM TECP and 10 mM D-Desthiobiotin) Further purifications were performed by size exclusion chromatography with the same buffer as ZYG11B-EloBC complex. The purified homogenous CUL2/Rbx-1 complex was verified by SDS-PAGE. Samples were concentrated and stored at -80 °C.

The purified ZYG11B-EloBC complex and CUL2/Rbx-1 complex were mixed with a ratio of 1:1 and left on ice for 30 minutes. The pentamer complex (ZBCC) was purified by size exclusion chromatography column (Superose 6 Increase 10/300 GL, GE Healthcare). Purified ZBCC complex were concentrated and stored at -80 °C for later usage.

Full-length Cyclin B1 protein and its fusion protein with NLRP1 N degrons were cloned into a pET-28a vector with an N-terminus 6xHis tag. The target protein was expressed and purified from *E. coli,* as described previously^43^. The N-degron was exposed by digestion with TEV protease with further purification by anion exchange chromatography (HiTrap Q HP column, Cytiva). The human CUL5 and Rbx2 protein complex was expressed and purified as described^44^. SARS-CoV ORF10 protein (2-38) and FITC labeled peptide of different Gly/N degron synthesized by Peptide 2.0 Inc.

Purified ZBC complex was mixed with ORF10 peptide in a molar ratio 1:10, yielding ZBC ORF10 complex for Cryo-EM analysis.

### MBP Pulldown assays

GSERRVLRQL and GSERRVLRQLPDTSGRRWRE were appended to the N terminus of MBP and constructed into pET-28a vector containing a 6x His tag followed by a TEV cleavage site. The fusion protein was purified using Ni-NTA agarose resin and digested by TEV protease. The digested protein was reloaded to Ni-NTA resin to remove TEV and 6xHis tag, followed by a sizing exclusion chromatography for further purification. The purified protein bears a Gly/N degron of GSERRVLRQL and GSERRVLRQLPDTSGRRWRE polypeptide overhang of MBP. Approximately 100 µg of MBP N-degron protein were immobilized with 50 µl amylose resin (NEB), followed by incubation with a molar ratio of 1:1 of target prey protein for 1 hr. at 4 C. After three times washing, the pull-downed sample was eluted with 100 µl sizing buffer adding 20 mM of Maltose. Samples were analyzed using SDS-PAGE with Coomassie Blue staining.

### Fluorescence Polarization assays

Different concentrations of purified ZYG11B/EloB/C complex were diluted with the Sizing buffer for protein purification (20mM HEPES pH 7.5, 300 mM NaCl, 0.5mM TCEP). FlTC-labeled peptides (Peptide 2.0) were dissolved in 100% DMSO adjusted to concentration of 10 mM for stock and diluted with Sizing buffer to 1 μM before conducting experiments. The binding assays were performed at 25° in Greiner Bio-One 384-Well low volume low binding plates. Protein and peptides were incubated for 20 minutes before measuring polarization on a SpectraMax plate reader (Molecular Dynamics).

Equilibrium dissociation constants (Kd) were fit to the Hill equation one site binding.

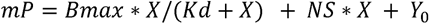

Where X represents protein concentration, Y_0_ is the background mP value of the peptides alone, Bmax is the mP value at saturation, and NS is the slope of nonlinear regression in mP value divided by X units.

### Cell culture and in vivo Co-IP assays

HEK 293T cells were maintained in DMEM (10% FBS, 1% pen/strep) at 37C with 5% CO2. The cells were transfected with 5 μg total DNA containing STREP-tagged NLRP1 or FLAG-tagged ZYG11B and its mutants. After 48 hours, cells were collected, lysed in Lysis Buffer (Strep lysis buffer:50 mM Tris-HCl pH 7.5, 150 mM NaCl, 2 mM MgCl_2_, 0.5% Triton X-100, 0.2% Na-deoxycholate, Roche protease inhibitor cocktail; Flag lysis buffer: 50 mM Tris-HCl pH 8.0, 150 mM NaCl, 1.5 mM MgCl2, 0.2 mM EDTA, 1% IGEPAL® CA-630, Roche protease inhibitor cocktail) and rocked for 30 minutes at 4[. Lysate was cleared and incubated with Strep-Tactin® Superflow® resin (IBA LifeScience) or magnetic M2-FLAG resin slurry (Sigma) at 4[. The resin was washed with WASH buffer (Strep WASH buffer:50 mM Tris-HCl pH 7.5, 150 mM NaCl, 2 mM MgCl2, 0.5% Triton X-100; Flag WASH buffer: 50 mM Tris-HCl pH 8.0, 150 mM NaCl, 1.5 mM MgCl2, 0.2mM EDTA, 0.1% IGEPAL® CA-630) and eluted with 1x STREP elution buffer (IBA-GmbH) diluted in Strep WASH buffer or FLAG elution buffer (200 μg/mL 3x FLAG peptide, diluted in Flag WASH buffer), shaking at RT for 30 minutes. The eluate was mixed 1:1 with 2x Laemmli sample buffer, boiled, and loaded on SDS-PAGE gel. Protein samples were transferred to PVDF membrane by using semi-dry transfer system. Target proteins were detected with specific antibodies.

### Inflammasome assays

HEK293, AAN2, and ZYG11B knockout cells were seeded the day prior to transfection in a 24-well plate (Genesee, El Cajon, CA) with 500 µl complete media containing DMEM (Gibco, Carlsbad, CA), 10% FBS (Peak Serum, Wellington, CO), and appropriate antibiotics (Gibco, Carlsbad, CA). Cells were transiently transfected with 500 ng of total DNA and 1.5 µl of Transit X2 (Mirus Bio, Madison, WI) following the manufacturer’s protocol. NLRP1 inflammasome activation assays were performed similarly to those previously described ^33^. Briefly, 5 ng ASC, 100 ng CASP1, 50 ng IL-1β-V5, and 8 ng of pQCXIP-NLRP1-TEV-Myc constructs were co-transfected with 250ng of HA-tagged TEV protease, 100ng HA-tagged HRV 3Cprotease construct or empty vector (pQCXIP). For experiments investigating SARS-CoV-2 ORF10 inhibition of NLRP1 activation 5 ng ASC, 100 ng CASP1, 50 ng IL-1β-V5, and 8 ng of pCDNA-4TO-2*strep-hNLRP1 or pCDNA-4TO-2*strep-hNLRP1-E133K constructs were co-transfected in HEK293 cells with 100 ng HA-tagged HRV 3Cprotease, HA-tagged HRV 3Cprotease and pCDNA-4TO-2*strep -SARS-CoV-2-ORF10, or pQCXIP. Twenty-two hours post-transfection, cells were washed with 1X PBS, harvested, and lysed in 1x NuPAGE LDS sample buffer (Invitrogen, Carlsbad, CA) containing 5% β-mercaptoethanol (Fisher Scientific, Pittsburg, PA) at 98C for 10 min. The lysed samples were loaded into 4-12% Bis-Tris SDS-PAGE gel with 1X MES buffer and wet transferred onto a nitrocellulose membrane (Life Technologies, San Diego, CA). Membranes were blocked with PBS-T containing 5% bovine serum albumin (BSA), followed by incubation with primary antibodies for V5 (IL-1β), Myc (NLRP1), or Strep (NLRP1 and NLRP1-E133K), HA (HRV 3C protease) or GAPDH. Membranes were rinsed three times for 5 min in PBS-T and then incubated with the appropriate HRP-conjugated secondary antibodies. Membranes were rinsed again three times in PBS-T and developed with SuperSignal West Pico PLUS Chemiluminescent Substrate (Thermo Fisher Scientific, Carlsbad, CA). Relative densitometry units (p17-pro-IL-1β) were quantified using FIJI.

### Electron-microscopy data acquisition

For negative stain, 2.5 µl of sample was applied to glow-discharged EM grids covered by a thin layer of carbon film (TED Pella, Inc) and stained with 0.75% (w/v) uranyl formate solution. Negative stain grids were imaged on a Tecnai T12 microscope (Thermo Fisher Scientific) operated at 120kV with a camera UltraScan4000(Gatan Inc). Images were recorded at a magnification of x52,000, in a 2.23 -Å pixel size on the specimen. Defocus was set to −1.5 µm. For Cryo-EM, 3□µl of sample (∼1.0□mg/ml) was applied to quanlifoil grids The grids were blotted by filter paper and plunge-frozen in liquid ethane using a Mark IV Vitrobot (Thermo Fisher Scientific) with blotting times of 4–7 □s at room temperature and over 90% humidity. Cryo-EM datasets were collected using Serial EM under 300kV on Titan Krios microscopes equipped with Field Emission source and K3 camera (Gatan Inc.)

### Image processing and model building

For all Cryo-EM datasets, movies were motion-corrected by MotionCor2. Motion-corrected sums without dose weighting were used for defocus estimation by using Gctf. Motion-corrected sums with dose weighting were used for all other image processing. All datasets were processed similarly. In summary, particle picking,2D class averages, initial model generation, and auto-refinement by following the workflow in Relion 3.1 or 4.0^45^. Final maps were refined, reconstructed, and sharpened in Phenix. The resolution was estimated by the FSC[=[0.143 criterion. The atomic model of CUL2-Rbx1-Elongin-B/C(PDB:5N4W ) was used as a starting model, followed by manually adjusting in COOT to fit the density map, ZYG11B and N degron of NLRP1 were manually built and adjusted in COOT with the help of RossetaCommons model prediction. The final model was refined with Phenix and adjusted in COOT repeat until Ramachandran validation was satisfied^46^. UCSF Chimera X and Pymol were used to prepare images^47^.

## Notes

### Competing Interest Statement

The authors have declared no competing interest.

## Uncategorized References

1 Harper, J. W. & Schulman, B. A. Cullin-RING Ubiquitin Ligase Regulatory Circuits: A Quarter Century Beyond the F-Box Hypothesis. Annu Rev Biochem 90, 403–429 (2021). 10.1146/annurev-biochem-090120-013613

2 Timms, R. T. et al. A glycine-specific N-degron pathway mediates the quality control of protein N-myristoylation. Science 365 (2019). 10.1126/science.aaw4912

3 Yan, X. et al. Molecular basis for recognition of Gly/N-degrons by CRL2(ZYG11B) and CRL2(ZER1). Mol Cell 81, 3262–3274 e3263 (2021). 10.1016/j.molcel.2021.06.010

4 Li, Y. et al. CRL2(ZER1/ZYG11B) recognizes small N-terminal residues for degradation. Nat Commun 13, 7636 (2022). 10.1038/s41467-022-35169-6

5 Balachandran, R. S. et al. The ubiquitin ligase CRL2ZYG11 targets cyclin B1 for degradation in a conserved pathway that facilitates mitotic slippage. J Cell Biol 215, 151–166 (2016). 10.1083/jcb.201601083

6 Zhang, B. et al. Structural insights into ORF10 recognition by ZYG11B. Biochem Biophys Res Commun 616, 14–18 (2022). 10.1016/j.bbrc.2022.05.069

7 Robinson, K. S. et al. Enteroviral 3C protease activates the human NLRP1 inflammasome in airway epithelia. Science 370 (2020). 10.1126/science.aay2002

8 Levinsohn, J. L. et al. Anthrax lethal factor cleavage of Nlrp1 is required for activation of the inflammasome. PLoS Pathog 8, e1002638 (2012). 10.1371/journal.ppat.1002638

9 Chavarria-Smith, J. & Vance, R. E. Direct proteolytic cleavage of NLRP1B is necessary and sufficient for inflammasome activation by anthrax lethal factor. PLoS Pathog 9, e1003452 (2013). 10.1371/journal.ppat.1003452

10 Planes, R. et al. Human NLRP1 is a sensor of pathogenic coronavirus 3CL proteases in lung epithelial cells. Mol Cell 82, 2385–2400 e2389 (2022). 10.1016/j.molcel.2022.04.033

11 Chui, A. J. et al. N-terminal degradation activates the NLRP1B inflammasome. Science 364, 82–85 (2019). 10.1126/science.aau1208

12 Sandstrom, A. et al. Functional degradation: A mechanism of NLRP1 inflammasome activation by diverse pathogen enzymes. Science 364 (2019). 10.1126/science.aau1330

13 Mitchell, P. S., Sandstrom, A. & Vance, R. E. The NLRP1 inflammasome: new mechanistic insights and unresolved mysteries. Curr Opin Immunol 60, 37–45 (2019). 10.1016/j.coi.2019.04.015

14 Hollingsworth, L. R. et al. DPP9 sequesters the C terminus of NLRP1 to repress inflammasome activation. Nature 592, 778–783 (2021). 10.1038/s41586-021-03350-4

15 Huang, M. et al. Structural and biochemical mechanisms of NLRP1 inhibition by DPP9. Nature 592, 773–777 (2021). 10.1038/s41586-021-03320-w

16 Castro, L. K. & Daugherty, M. D. Tripping the wire: sensing of viral protease activity by CARD8 and NLRP1 inflammasomes. Curr Opin Immunol 83, 102354 (2023). 10.1016/j.coi.2023.102354

17 Gong, Q. et al. Structural basis for distinct inflammasome complex assembly by human NLRP1 and CARD8. Nat Commun 12, 188 (2021). 10.1038/s41467-020-20319-5

18 Taabazuing, C. Y., Griswold, A. R. & Bachovchin, D. A. The NLRP1 and CARD8 inflammasomes. Immunol Rev 297, 13–25 (2020). 10.1111/imr.12884

19 Tsu, B. V. et al. Host-specific sensing of coronaviruses and picornaviruses by the CARD8 inflammasome. PLoS Biol 21, e3002144 (2023). 10.1371/journal.pbio.3002144

20 Kulsuptrakul, J., Turcotte, E. A., Emerman, M. & Mitchell, P. S. A human-specific motif facilitates CARD8 inflammasome activation after HIV-1 infection. Elife 12 (2023). 10.7554/eLife.84108

21 Wang, Q. et al. CARD8 is an inflammasome sensor for HIV-1 protease activity. Science 371 (2021). 10.1126/science.abe1707

22 Clark, K. M., Pal, P., Kim, J. G., Wang, Q. & Shan, L. The CARD8 inflammasome in HIV infection. Adv Immunol 157, 59–100 (2023). 10.1016/bs.ai.2022.11.001

23 Soucy, T. A. et al. An inhibitor of NEDD8-activating enzyme as a new approach to treat cancer. Nature 458, 732–736 (2009). 10.1038/nature07884

24 Zhang, J. et al. ZYG11B potentiates the antiviral innate immune response by enhancing cGAS-DNA binding and condensation. Cell Rep 42, 112278 (2023). 10.1016/j.celrep.2023.112278

25 Gordon, D. E. et al. A SARS-CoV-2 protein interaction map reveals targets for drug repurposing. Nature 583, 459–468 (2020). 10.1038/s41586-020-2286-9

26 Li, X. et al. SARS-CoV-2 ORF10 suppresses the antiviral innate immune response by degrading MAVS through mitophagy. Cell Mol Immunol 19, 67–78 (2022). 10.1038/s41423-021-00807-4

27 Stebbins, C. E., Kaelin, W. G., Jr. & Pavletich, N. P. Structure of the VHL-ElonginC-ElonginB complex: implications for VHL tumor suppressor function. Science 284, 455–461 (1999). 10.1126/science.284.5413.455

28 Huber, A. H., Nelson, W. J. & Weis, W. I. Three-dimensional structure of the armadillo repeat region of beta-catenin. Cell 90, 871–882 (1997). 10.1016/s0092-8674(00)80352-9

29 Tewari, R., Bailes, E., Bunting, K. A. & Coates, J. C. Armadillo-repeat protein functions: questions for little creatures. Trends Cell Biol 20, 470–481 (2010). 10.1016/j.tcb.2010.05.003

30 Nguyen, H. C., Yang, H., Fribourgh, J. L., Wolfe, L. S. & Xiong, Y. Insights into Cullin-RING E3 ubiquitin ligase recruitment: structure of the VHL-EloBC-Cul2 complex. Structure 23, 441–449 (2015). 10.1016/j.str.2014.12.014

31 Kamura, T. et al. VHL-box and SOCS-box domains determine binding specificity for Cul2-Rbx1 and Cul5-Rbx2 modules of ubiquitin ligases. Genes Dev 18, 3055–3065 (2004). 10.1101/gad.1252404

32 Zhou, H., Zaher, M. S., Walter, J. C. & Brown, A. Structure of CRL2Lrr1, the E3 ubiquitin ligase that promotes DNA replication termination in vertebrates. Nucleic Acids Res 49, 13194–13206 (2021). 10.1093/nar/gkab1174

33 Tsu, B. V. et al. Diverse viral proteases activate the NLRP1 inflammasome. Elife 10 (2021). 10.7554/eLife.60609

34 Kim, D. et al. The Architecture of SARS-CoV-2 Transcriptome. Cell 181, 914–921 e910 (2020). 10.1016/j.cell.2020.04.011

35 Davidson, A. D. et al. Characterisation of the transcriptome and proteome of SARS-CoV-2 reveals a cell passage induced in-frame deletion of the furin-like cleavage site from the spike glycoprotein. Genome Med 12, 68 (2020). 10.1186/s13073-020-00763-0

36 Finkel, Y. et al. The coding capacity of SARS-CoV-2. Nature 589, 125–130 (2021). 10.1038/s41586-020-2739-1

37 Mena, E. L. et al. ORF10-Cullin-2-ZYG11B complex is not required for SARS-CoV-2 infection. Proc Natl Acad Sci U S A 118 (2021). 10.1073/pnas.2023157118

38 Chen, Z. et al. Profiling of SARS-CoV-2 Subgenomic RNAs in Clinical Specimens. Microbiol Spectr 10, e0018222 (2022). 10.1128/spectrum.00182-22

39 Wang, L. et al. SARS-CoV-2 ORF10 impairs cilia by enhancing CUL2ZYG11B activity. J Cell Biol 221 (2022). 10.1083/jcb.202108015

40 Gu, S., et al. Recombinant SARS-CoV-2 lacking initiating and internal methionine codons within ORF10 is attenuated in vivo. bioRxiv (2023). 10.1101/2023.08.04.551973

41 Haltom, J., et al. SARS-CoV-2 Orphan Gene ORF10 Contributes to More Severe COVID-19 Disease. medRxiv (2023). 10.1101/2023.11.27.23298847

42 Bekes, M., Langley, D. R. & Crews, C. M. PROTAC targeted protein degraders: the past is prologue. Nat Rev Drug Discov 21, 181–200 (2022). 10.1038/s41573-021-00371-6

43 Hagting, A., Jackman, M., Simpson, K. & Pines, J. Translocation of cyclin B1 to the nucleus at prophase requires a phosphorylation-dependent nuclear import signal. Curr Biol 9, 680–689 (1999). 10.1016/s0960-9822(99)80308-x

44 Jager, S. et al. Vif hijacks CBF-beta to degrade APOBEC3G and promote HIV-1 infection. Nature 481, 371–375 (2011). 10.1038/nature10693

45 Scheres, S. H. RELION: implementation of a Bayesian approach to cryo-EM structure determination. J Struct Biol 180, 519–530 (2012). 10.1016/j.jsb.2012.09.006

46 Emsley, P., Lohkamp, B., Scott, W. G. & Cowtan, K. Features and development of Coot. Acta Crystallogr D Biol Crystallogr 66, 486–501 (2010). 10.1107/S0907444910007493

47 Meng, E. C. et al. UCSF ChimeraX: Tools for structure building and analysis. Protein Sci 32, e4792 (2023). 10.1002/pro.4792

