## Supplementary figures and images for "Structure of the E3 ligase CRL2-ZYG11B with substrates reveals the molecular basis for N-degron recognition and ubiquitination"

### Extended_figure 1

#### Extended Data Figure 1

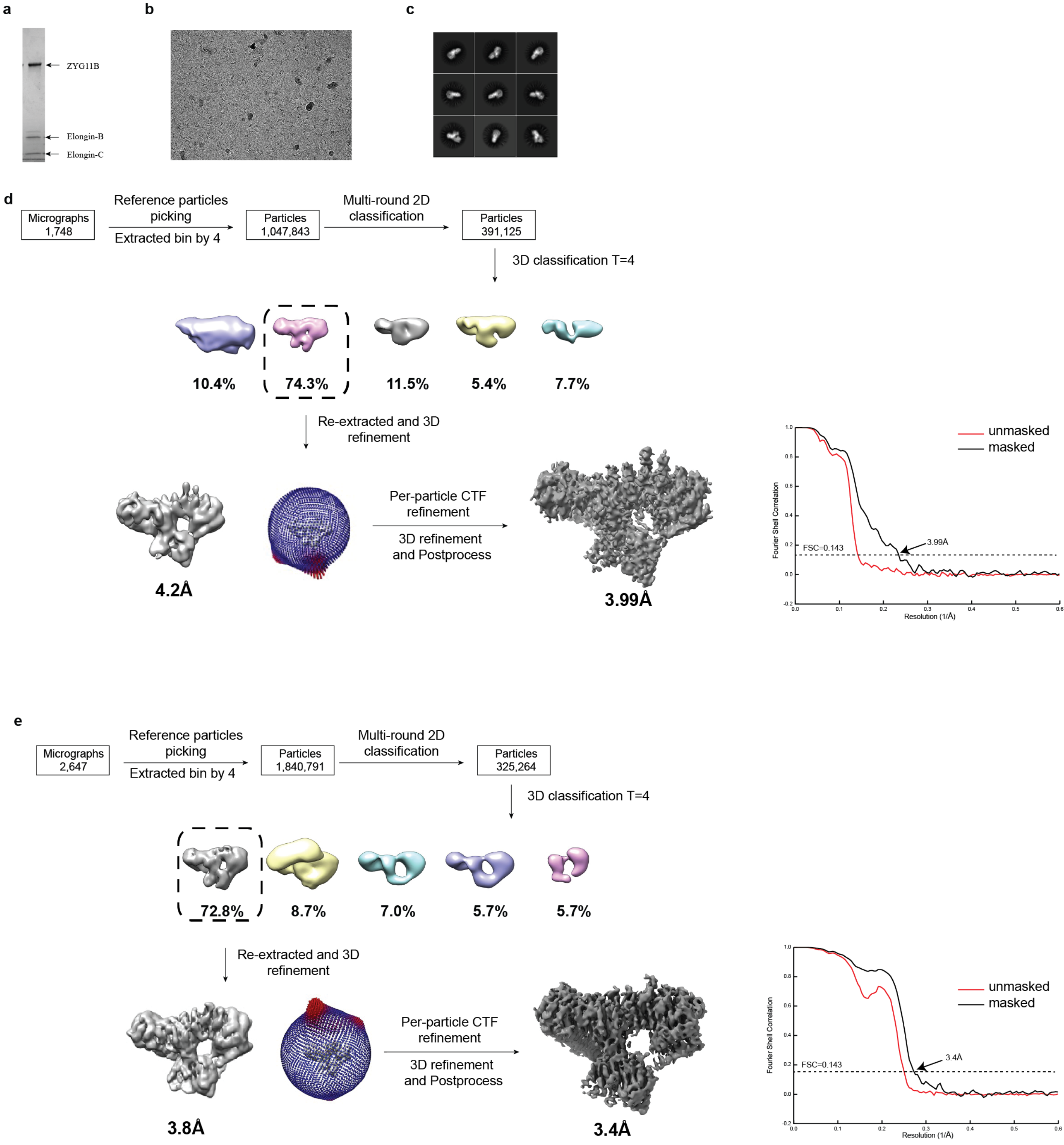

### Extended_figure 2

**Extended Data Figure 2**

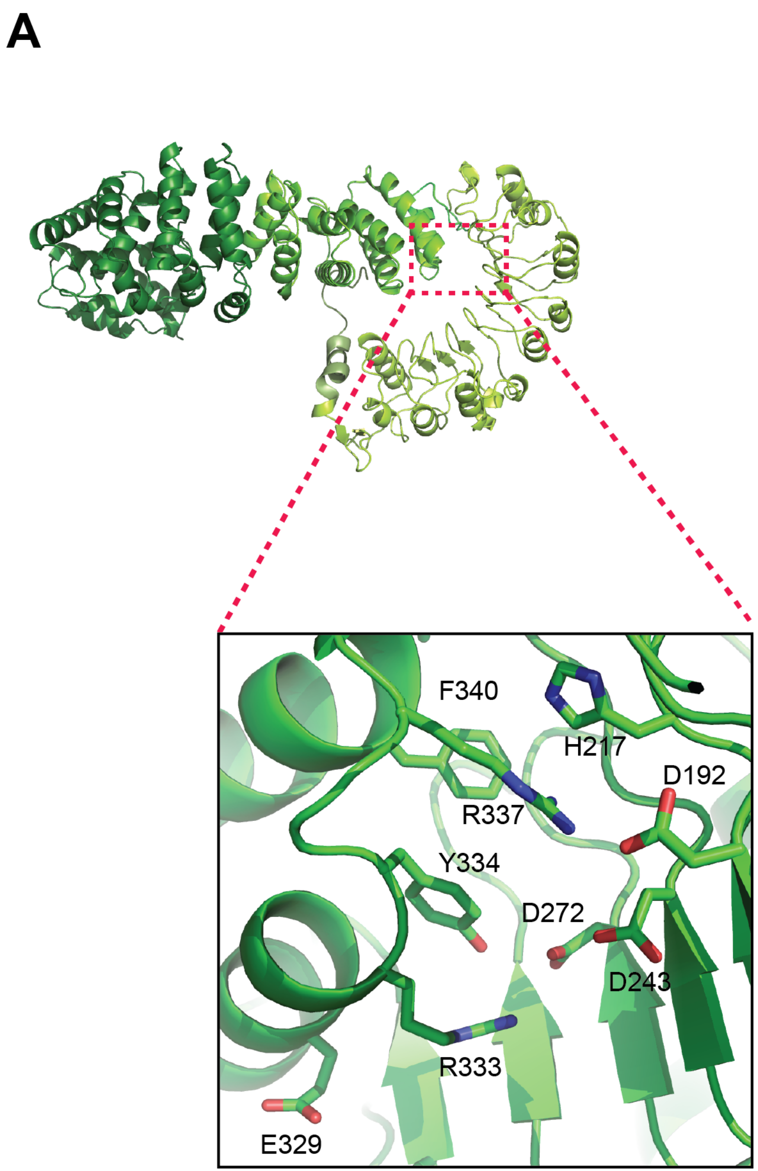


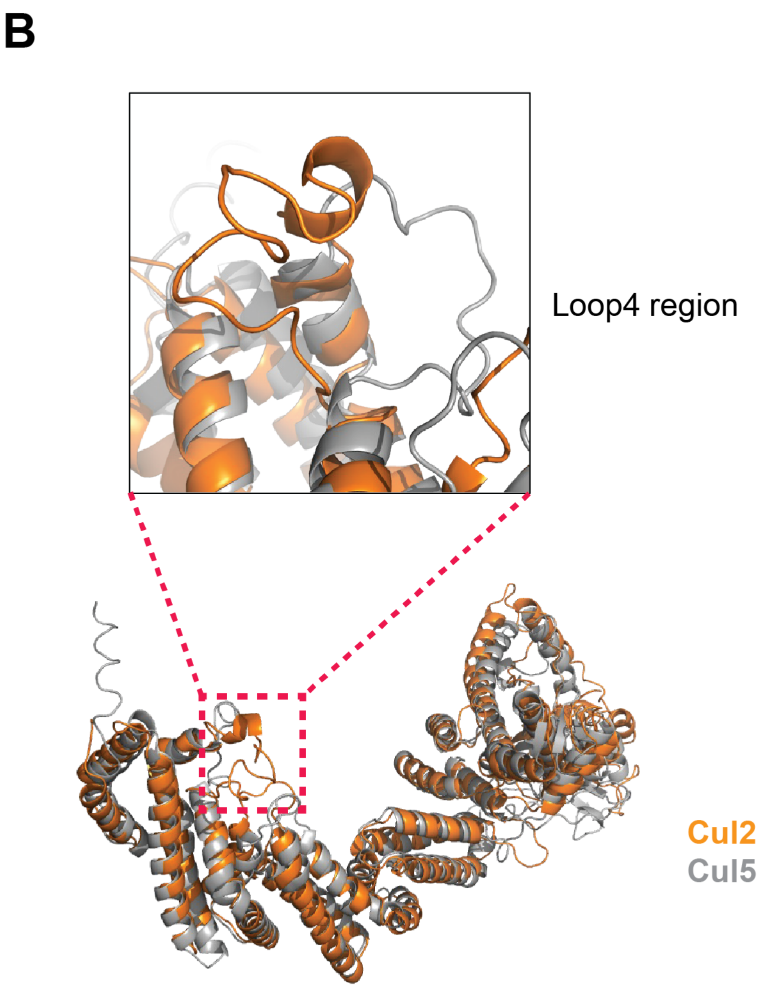

### Extended_figure 3

#### Extended Data Figure 3

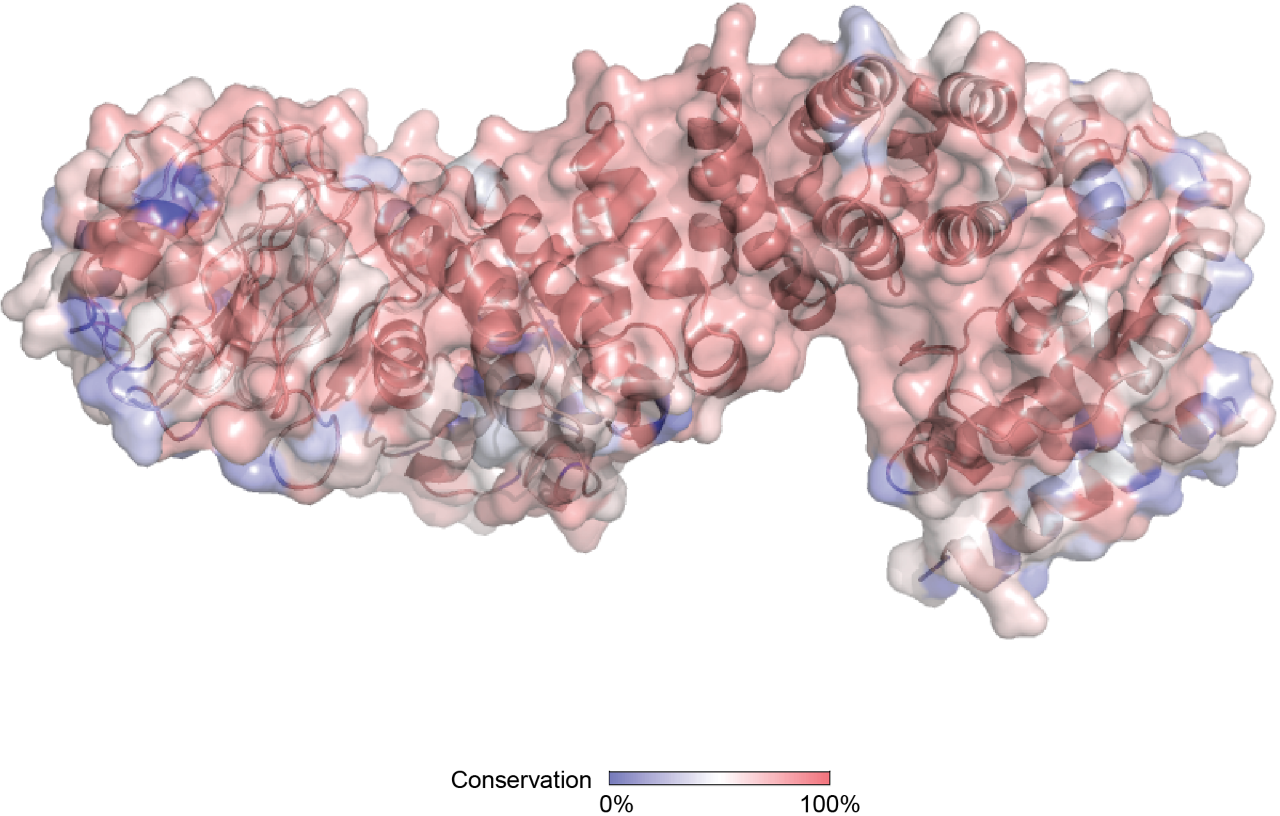


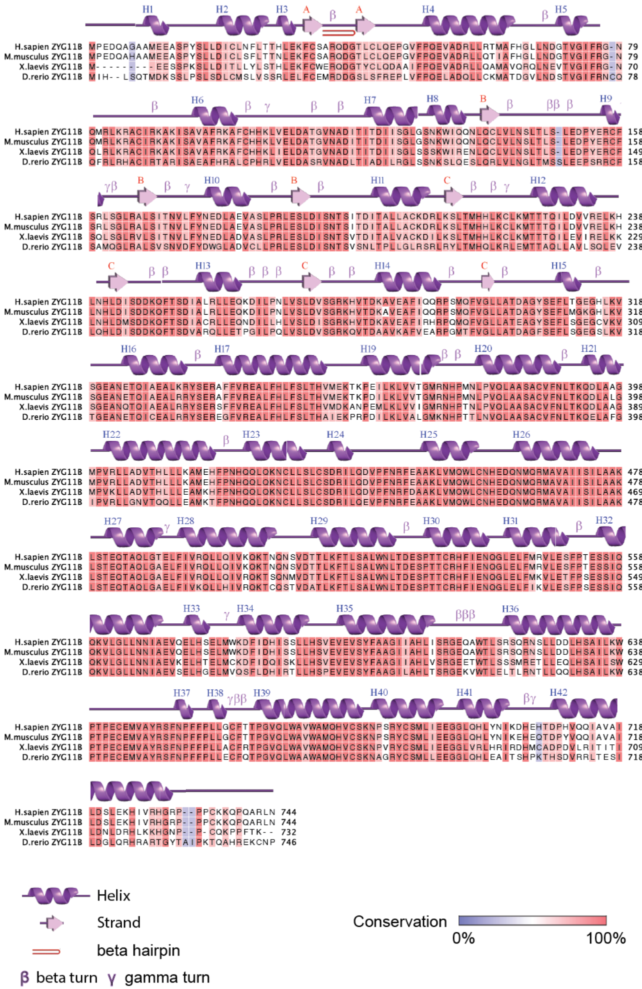

### Extended_figure 4

#### Extended Data Figure 4

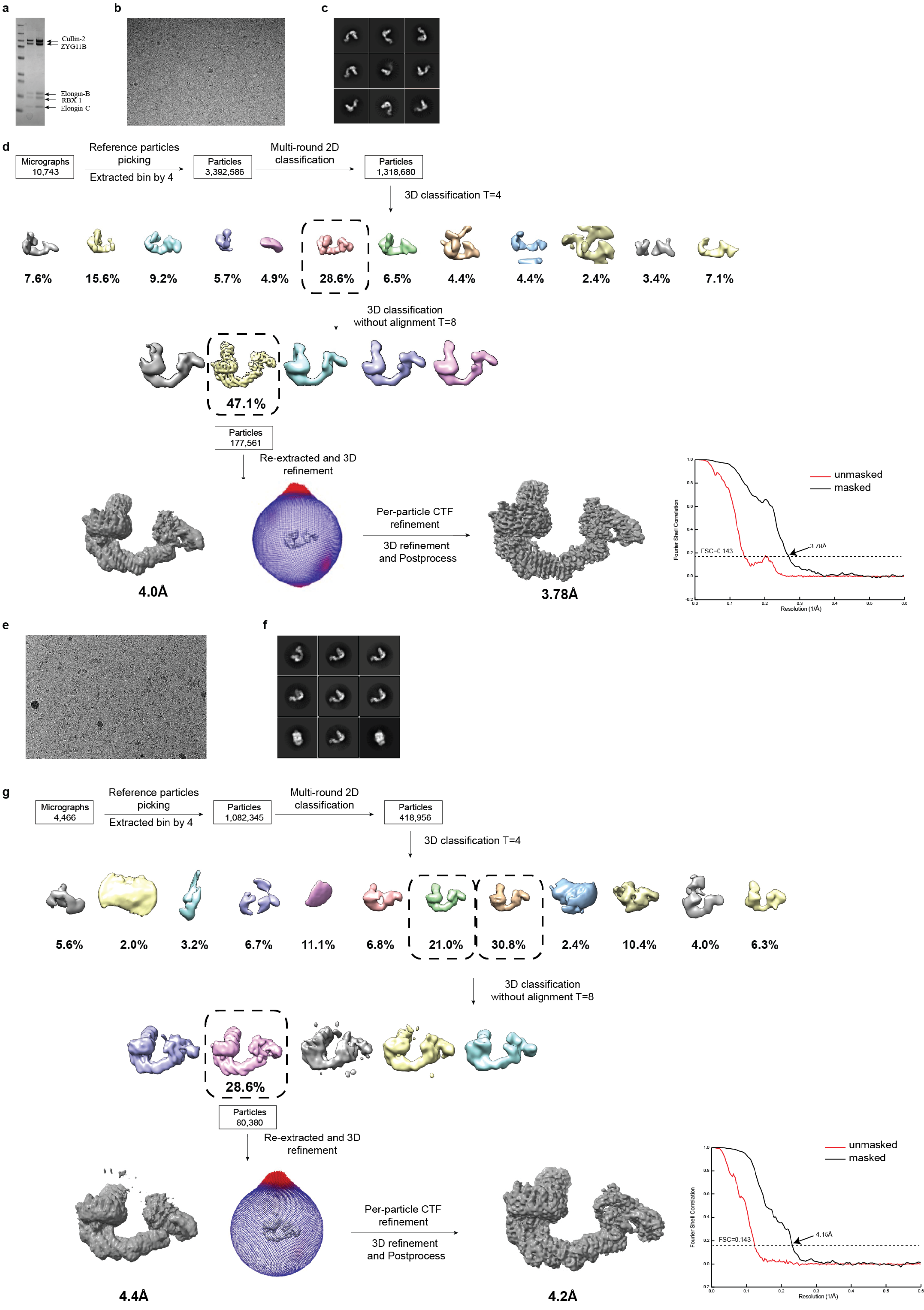

### Extended_figure 5

#### Extended Data Figure 5

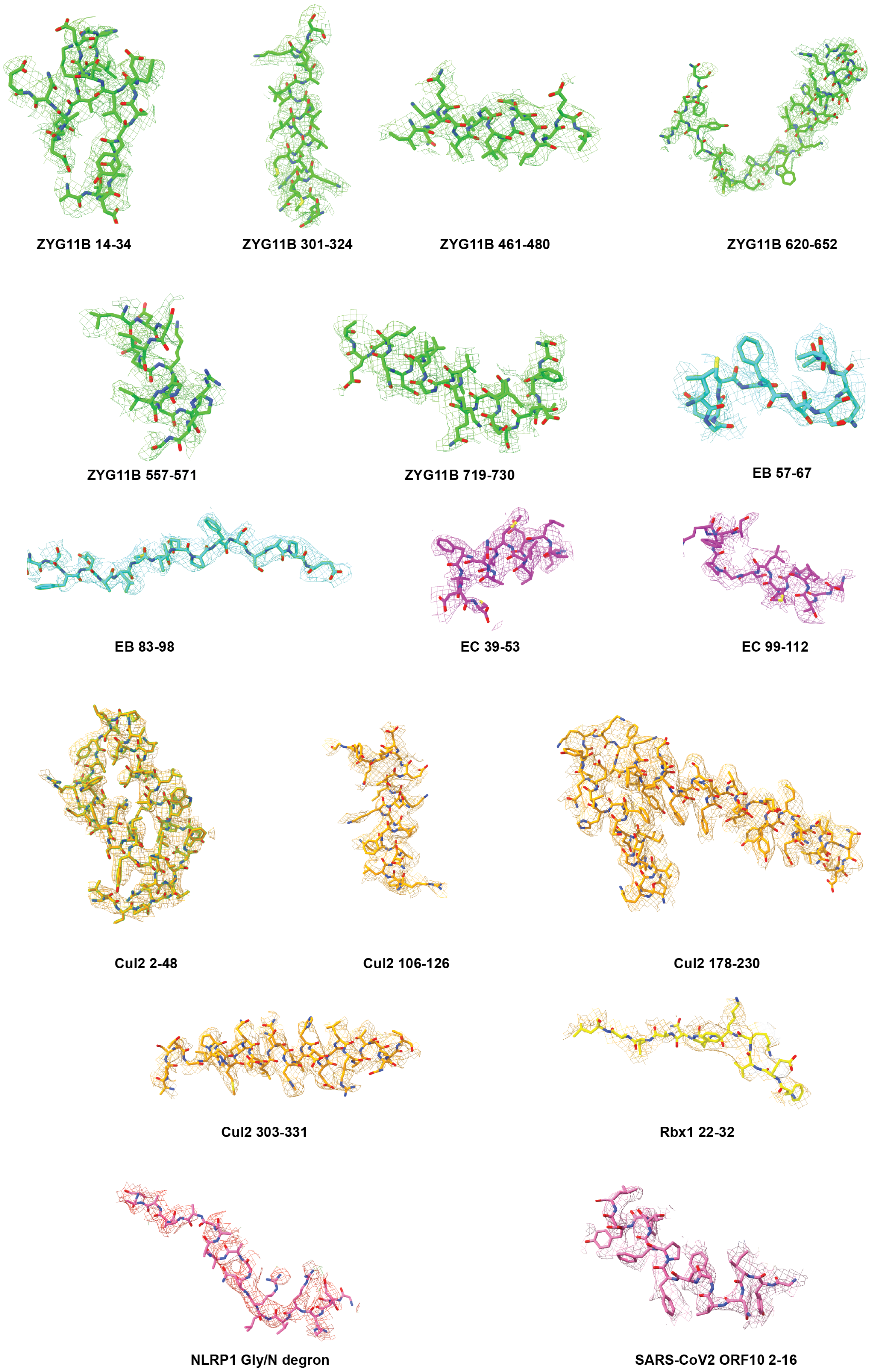

### Extended_figure 6

#### Extended Data Figure 6

A

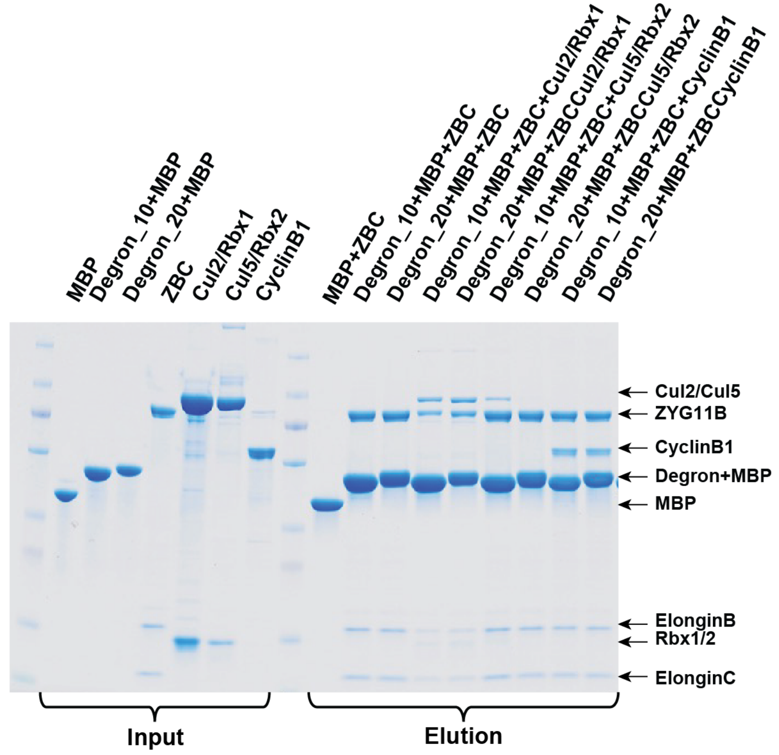


B

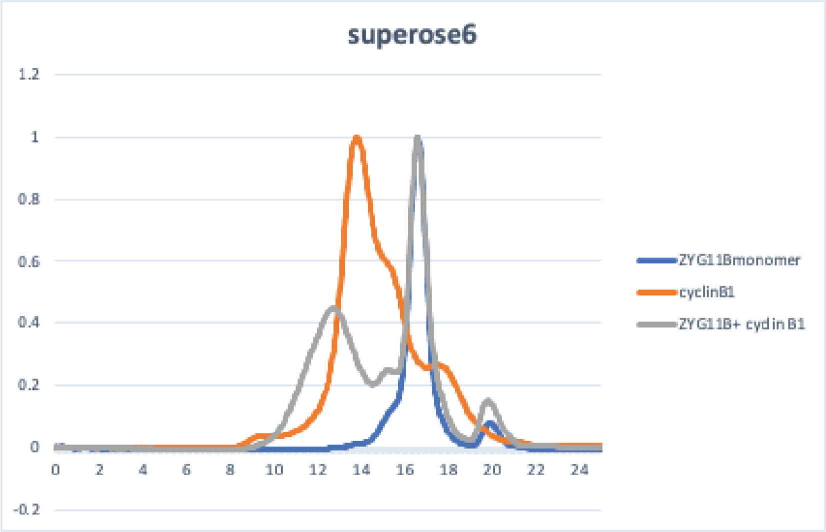


C

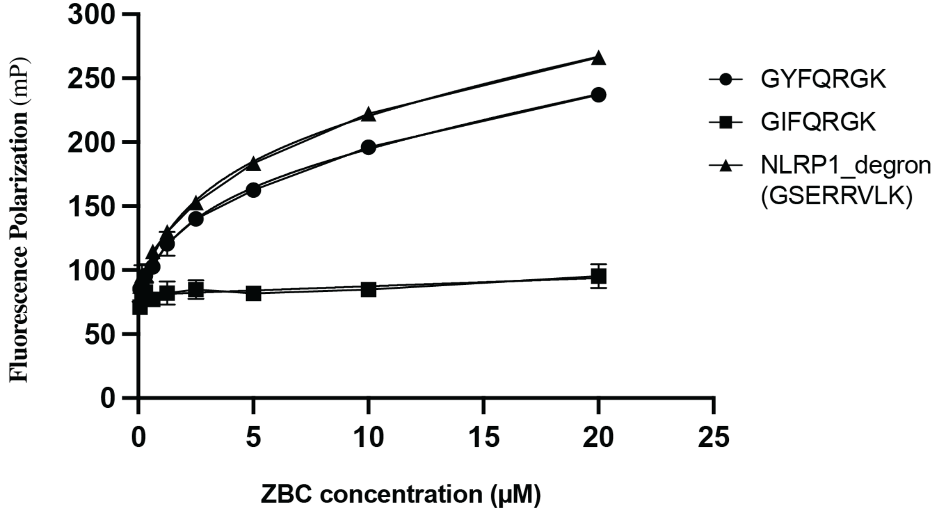


| Peptides | Kd(μM) |
|---|---|
| GYFQRGK | 2.5±0.1 |
| GIFQRGK | N.D |
| NLRP1_degron (GSERRVLK) | 2.9±0.3 |

### Extended_figure 7

#### Extended Data Figure 7

A

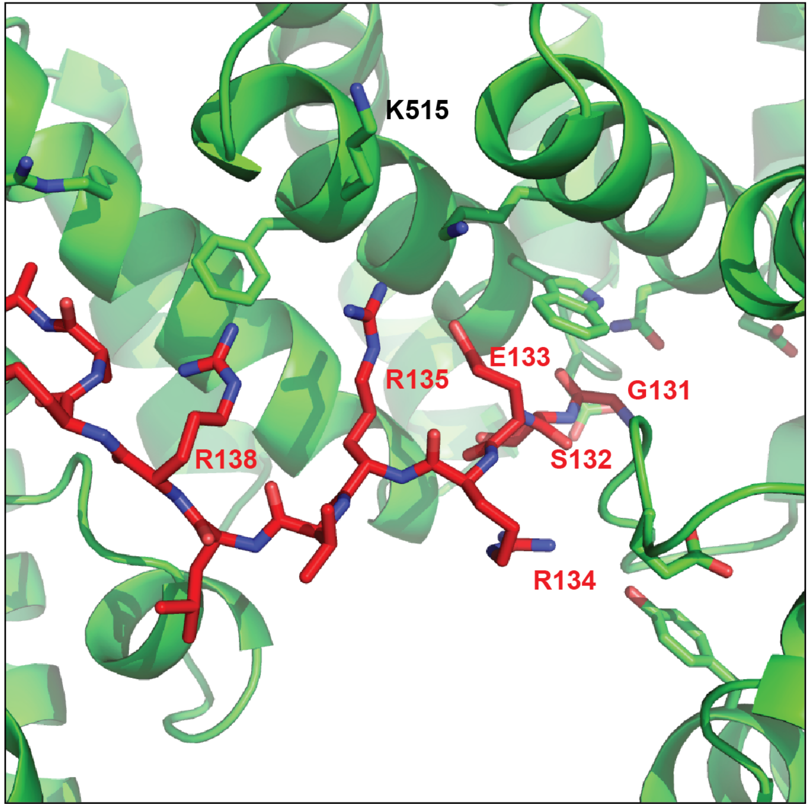


B

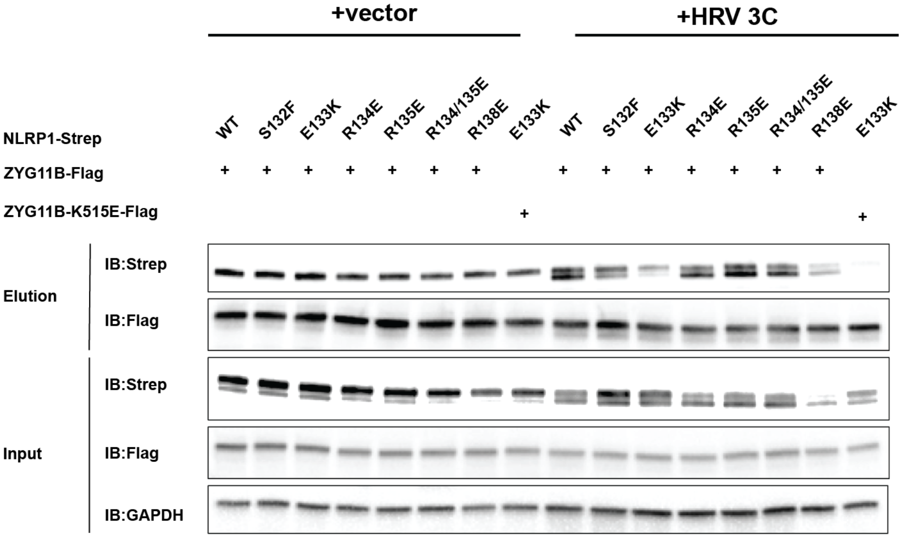
