## Supplementary material for "Structure of the E3 ligase CRL2-ZYG11B with substrates reveals the molecular basis for N-degron recognition and ubiquitination": table S1

### Cryo-EM data collection, refinement and validation statistics

| | ZYG11B/ElonginB/C (EMDB-44588) (PDB 9BID) | ZYG11B/ElonginB/C plus Orf10 peptide (EMDB-44589) (PDB 9BIE) | ZYG11B/ElonginB/C/ Cullin-2/Rbx1 (EMDB- 44630) (PDB 9BJ8) | ZYG11B/ElonginB/C/ Cullin-2/Rbx-1 +CyclinB1-NLRP1-Gly/N degron (EMDB- 44631) (PDB 9BJ9) |
|---|---|---|---|---|
| **Data collection and processing** | | | | |
| Magnification | 105000 | 105000 | 105000 | 105000 |
| Voltage (kV) | 300 | 300 | 300 | 300 |
| Electron exposure (e–/Å$^2$) | 60 | 60 | 60 | 60 |
| Defocus range (μm) | -1.0 to -2.5 | -1.0 to -2.5 | -0.8 to -2.2 | -0.8 to -2.2 |
| Pixel size (Å) | 0.835 | 0.835 | 0.835 | 0.835 |
| Symmetry imposed | P1 | P1 | P1 | P1 |
| Initial particle images (no.) | 1047873 | 1840791 | 3392586 | 1082345 |
| Final particle images (no.) | 123976 | 325264 | 177561 | 80380 |
| Map resolution (Å) | 3.9 | 3.4 | 3.7 | 4.1 |
| FSC threshold | | | | |
| Map resolution range (Å) | 0.143 | 0.143 | 0.143 | 0.143 |
| **Refinement** | | | | |
| Initial model used (PDB code) | 4WQO | 4WQO | 5N4W | 5N4W |
| Model resolution (Å) | 3.99 | 3.5 | 3.78 | 4.13 |
| FSC threshold | | | | |
| Model resolution range (Å) | 3.9/4.1/8.3 | 2.8/3.2/6.6 | 3.1/3.8/6.6 | 3.5/4.1/7.0 |
| Map sharpening *B* factor (Å$^2$) | | | | |
| Model composition | | | | |
| Non-hydrogen atoms | 7249 | 7353 | 14069 | 14205 |
| Protein residues | 915 | 933 | 1746 | 1767 |
| Ligands | 0 | 0 | 3 | 3 |
| *B* factors (Å$^2$) | | | | |
| Protein | 9.46/195.16/44.58 | 23.46/126.16/62.05 | 36.22/540.47/133.75 | 50.1/468.04/159.74 |
| Ligand | | | | |
| R.m.s. deviations | | | | |
| Bond lengths (Å) | 0.818 | 0.777 | 0.757 | 0.795 |
| Bond angles (°) | 0.003 | 0.003 | 0.003 | 0.003 |
| Validation | | | | |
| MolProbity score | 2.09 | 1.88 | 1.93 | 2.00 |
| Clashscore | 17.18 | 14.94 | 12.23 | 14.50 |
| Poor rotamers (%) | 0.00 | 0.00 | 0.00 | 0.00 |
| Ramachandran plot | | | | |
| Favored (%) | 94.94 | 96.75 | 95.33 | 95.33 |
| Allowed (%) | 5.06 | 3.25 | 4.67 | 4.68 |
| Disallowed (%) | 0.00 | 0.00 | 0.00 | 0.00 |
